# Pigeon foot feathering reveals conserved limb identity networks

**DOI:** 10.1101/602987

**Authors:** Elena F. Boer, Hannah F. Van Hollebeke, Sungdae Park, Carlos R. Infante, Douglas B. Menke, Michael D. Shapiro

## Abstract

The tetrapod limb is a stunning example of evolutionary diversity, with dramatic variation not only among distantly related species, but also between the serially homologous forelimbs (FLs) and hindlimbs (HLs) within species. Despite this variation, highly conserved genetic and developmental programs underlie limb development and identity in all tetrapods, raising the question of how limb diversification is generated from a conserved toolkit. In some breeds of domestic pigeon, shifts in the expression of two conserved limb identity transcription factors, *PITX1* and *TBX5*, are associated with the formation of feathered HLs with partial FL identity. To determine how modulation of *PITX1* and *TBX5* expression affects downstream gene expression, we compared the transcriptomes of embryonic limb buds from pigeons with scaled and feathered HLs. We identified a set of differentially expressed genes enriched for genes encoding transcription factors, extracellular matrix proteins, and components of developmental signaling pathways with important roles in limb development. A subset of the genes that distinguish scaled and feathered HLs are also differentially expressed between FL and scaled HL buds in pigeons, pinpointing a set of gene expression changes downstream of *PITX1* and *TBX5* in the partial transformation from HL to FL identity. We extended our analyses by comparing pigeon limb bud transcriptomes to chicken, anole lizard, and mammalian datasets to identify deeply conserved *PITX1*- and *TBX5*-regulated components of the limb identity program. Our analyses reveal a suite of predominantly low-level gene expression changes that are conserved across amniotes to regulate the identity of morphologically distinct limbs.

**Summary statement:** In feather-footed pigeons, mutant alleles of *PITX1* and *TBX5* drive the partial redeployment of an evolutionarily conserved forelimb genetic program in the hindlimb.

## Introduction

The tetrapod limb is an extraordinary example of morphological variation. Dramatic evolutionary diversification of limb form and function has enabled animals to thrive in diverse terrestrial, fossorial, aquatic, and aerial environments. The limb is unusual in that phenotypic variation characterizes not only limbs of different species, but also the serially homologous forelimbs (FLs) and hindlimbs (HLs) within a species. Limb variation between FLs and HLs is arguably most striking in birds, in which divergent patterning of the muscles, tendons, and bones that form the limb is coupled with variation in the epidermal appendages that cover each limb type. For example, the elaborately feathered epidermis of the avian wing is dramatically different from the scaled epidermis that typically covers the feet.

Despite highly divergent adult limb morphologies among and within species, limb patterning and morphogenesis is orchestrated by deeply conserved genetic and developmental programs. As a result of decades of elegant descriptive and experimental studies in canonical model organisms (especially mouse and chicken), the cellular, genetic, and molecular regulators of limb development are relatively well defined (reviewed in (Petit et al., 2017, Tickle, 2015, Rabinowitz and Vokes, 2012, Duboc and Logan, 2011b, Towers and Tickle, 2009, Tickle, 2004)). These studies reveal that the integration of several key developmental signaling pathways – including Fibroblast Growth Factor (FGF), Bone Morphogenetic Protein (BMP), Wnt, Sonic Hedgehog (SHH), and Retinoic Acid (RA) pathways – is required for proper limb bud position, initiation, outgrowth, and patterning.

In contrast, FL and HL identity is associated with the reciprocal expression patterns of a smaller set of genes, including the transcription factors (TFs) *TBX5* (FL-specific expression), *TBX4* (HL-specific), and *PITX1* (HL-specific) (Simon et al., 1997, Logan et al., 1998, Szeto et al., 1996, Gibson-Brown et al., 1996, Shang et al., 1997). Over the past 20 years, a combination of misexpression and genetic studies have illuminated essential roles for each of these TFs in limb development. The paralogs *TBX5* and *TBX4* direct the respective initiation of FL and HL bud development through activation of the FGF and Wnt signaling pathways (Takeuchi et al., 2003). In mice, *Tbx5^-/-^* mutants fail to form a FL bud, while *Tbx4^-/-^* mutants form a HL bud that arrests early in development due to a loss of *FGF10* expression maintenance (Agarwal et al., 2003, Takeuchi et al., 1999, Rallis et al., 2003, Naiche and Papaioannou, 2003, Naiche and Papaioannou, 2007). Similarly, loss of *tbx5* function in zebrafish results in a failure to form the pectoral fin (FL equivalent) (Ahn et al., 2002, Garrity et al., 2002), and disruption of *tbx4* nuclear localization in the naturally occurring *pelvic finless* zebrafish strain is associated with arrested pelvic fin (HL equivalent) development (Don et al., 2016). *Tbx5* and *Tbx4* are not essential for development of limb-specific identities in mice (Hasson et al., 2007, Naiche and Papaioannou, 2007, Minguillon et al., 2005); however, misexpression of *TBX4* in the FL or *TBX5* in the HL of chick embryos causes a partial transformation in limb identity in a subset of embryos, including changes in digit number, limb flexure, and epidermal appendage type (Rodriguez-Esteban et al., 1999, Takeuchi et al., 1999). Although the basis for the discrepancies between mouse and chick experiments remains unclear, it is possible that *TBX5* and *TBX4* have subtly different functions in mammals and birds, which diverged more than 300 million years ago (Horton et al., 2008).

Unlike *TBX4* and *TBX5*, *PITX1* is not required for limb bud initiation or outgrowth, but instead regulates these processes indirectly via activation of *TBX4* (Minguillon et al., 2005, Szeto et al., 1999, Logan and Tabin, 1999, Duboc and Logan, 2011a). In addition to *TBX4*-dependent regulation of HL outgrowth, *PITX1* serves a second essential role in limb development via *TBX4*-independent regulation of HL identity (Duboc and Logan, 2011a). In *Pitx1^-/-^* mutant mice, HLs develop, but lack HL-specific morphological characteristics (Szeto et al., 1999). In both chick and mouse embryos, ectopic expression of *PITX1* in the developing FL bud causes a partial transformation to a more HL-like identity (DeLaurier et al., 2006, Logan and Tabin, 1999). Similarly, mutations at the human *PITX1* locus are associated with HL-like characteristics in the FL (Al-Qattan et al., 2013) and congenital HL defects (Gurnett et al., 2008). *PITX1* regulatory mutations have also been discovered in natural populations of three-spine stickleback fish, in which loss of *PITX1* gene expression contributes to the adaptive loss of pelvic (HL) structures (Shapiro et al., 2004, Chan et al., 2010).

Substantial research efforts over the past two decades have helped decipher the genetic networks regulated by *PITX1*, *TBX4*, and *TBX5* during limb development (reviewed in (Duboc and Logan, 2011b, Rabinowitz and Vokes, 2012, Tickle, 2015, Petit et al., 2017)). More recently, contemporary genomic approaches have opened new avenues for defining downstream targets in an unbiased manner. Particular progress has been made in defining the network downstream of *PITX1* by comparing limb buds from wild-type and *Pitx1^-/-^* mutant mice using massively parallel sequencing of both RNA transcripts to identify differentially-expressed genes and DNA regions captured by chromatin immunoprecipitation to locate putative regulatory regions (RNA-seq and ChIP-seq, respectively) (Nemec et al., 2017, Infante et al., 2013, Wang et al., 2018). These approaches have resulted in a more comprehensive view of *PITX1*-dependent gene networks (and could be used to similarly reveal *TBX5*- and *TBX4*-dependent networks). Despite these important advances, we still have a limited understanding of the transcriptomic changes that result from less dramatic (non-binary) shifts in gene expression that are associated with evolutionary diversification of limb morphology.

The domestic pigeon (*Columba livia*) is an outstanding model to study the evolution of genetic and developmental programs that underlie limb diversification. Pigeons display striking variation in HL morphology within a single species and are amenable to genetic crosses, genomic analyses, and embryonic studies (Shapiro et al., 2013, Shapiro and Domyan, 2013, Domyan and Shapiro, 2017). While most pigeons have scaled HLs, in some breeds scaled epidermis is replaced by skin with a range of feather morphologies (Figure 1A-C). Classical genetic experiments show that large feather “muffs” are caused by the synergistic effects of mutant alleles at two loci, *grouse* (*gr*) and *Slipper* (*Sl*), which independently produce smaller foot feathers (Doncaster, 1912, Wexelsen, 1934, Hollander, 1937, Levi, 1986). We recently showed that *gr* and *Sl* are *cis*-regulatory alleles of the limb identity genes *PITX1* and *TBX5*, respectively (Domyan et al., 2016), and proposed a model in which feathered feet are the result of a partial transformation from HL to FL identity. This genetically-encoded, partial transformation in identity provides a unique opportunity to determine how altered expression of conserved limb identity genes modifies the downstream HL identity transcriptional program.

**Figure 1.**
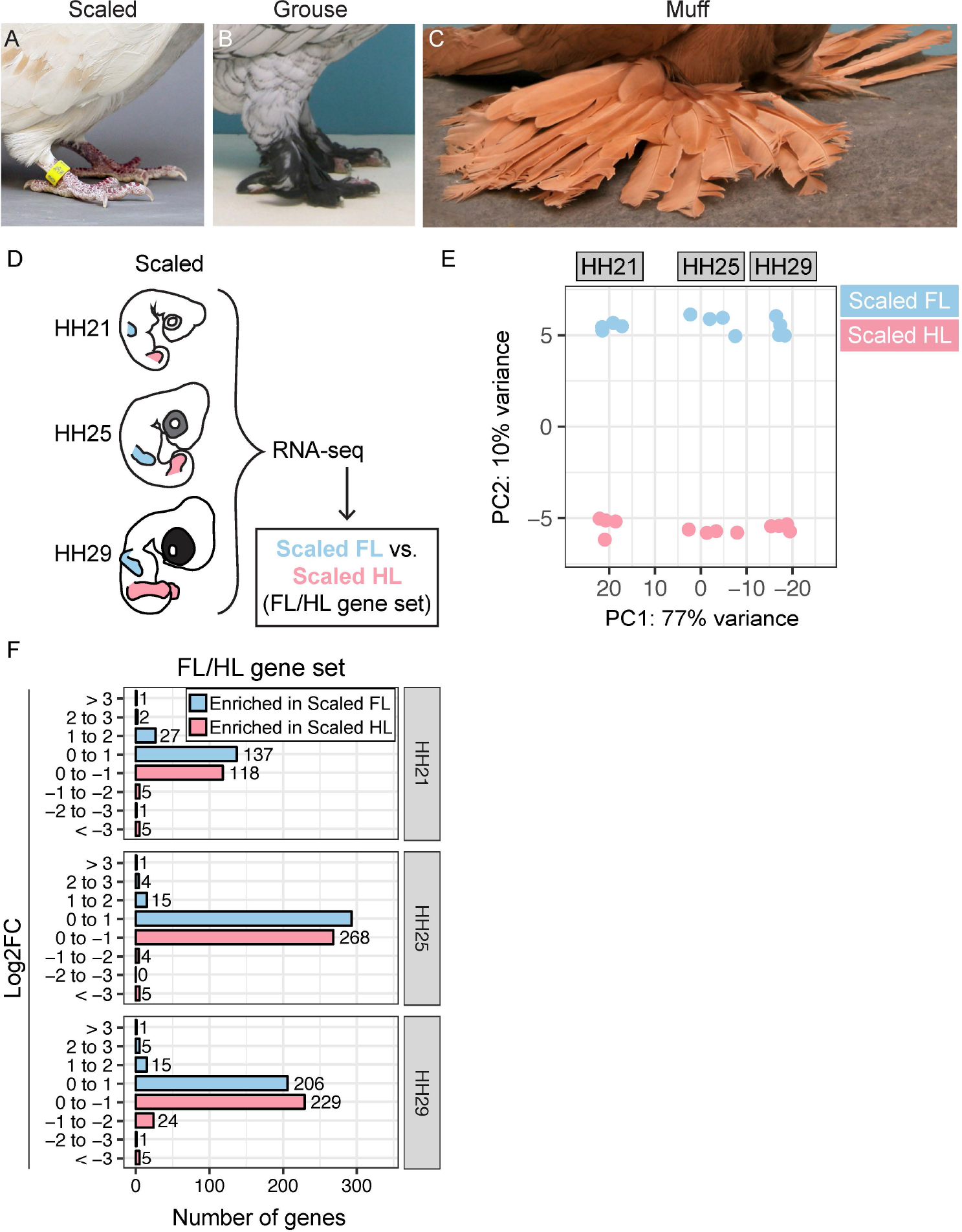
Comparative transcriptomics to identify gene expression changes associated with pigeon limb identity. (**A-C**) Hindlimb phenotypes in domestic pigeon breeds, including scaled (A), grouse (B), and muff (C). (**D**) Experimental approach to identify differentially expressed genes at three stages (HH21, HH25, and HH29) of pigeon FL vs. scaled HL development. (**E**) PCA based on top 500 most variable genes in FL and scaled HLs. PC1 (77% of variance) separates samples by stage, PC2 (10% of variance) separates samples by limb type. (**F**) Number of genes that show FL-enriched or scaled HL-enriched expression pattern at HH21, HH25, or HH29. Genes are grouped based on Log2FC. FL = forelimb; HL = hindlimb; HH = Hamburger-Hamilton stage.

The goal of this study is to identify genetic network changes that accompany shifts in limb identity. First, we identify FL- and HL-specific transcriptomic differences in pigeons. We then test for transcriptomic changes in pigeon HLs with partially altered limb identity attributable to changes in *PITX1* and *TBX5* regulation. Next, we compare our pigeon results with transcriptomic datasets from other amniotes to identify a set of genes whose expression typify FL and HL differences across broad phylogenetic distances. Finally, we find that a subset of the highly-conserved FL genetic program is also differentially expressed in mutant pigeon HLs with partial identity transformations.

## Results

### Gene expression changes associated with normal FL and HL development in pigeons

To identify differentially expressed transcripts associated with normal FL and HL development in pigeons, we performed RNA-seq on FL and HL buds from scale-footed pigeon embryos at three early stages of embryonic development (approximate pigeon equivalents of Hamburger-Hamilton (HH) stage 21, HH25, and HH29; (Hamburger and Hamilton, 1951)) (Figure 1D). Principal component analysis (PCA) based on the 500 most differentially expressed genes revealed that PC1 accounts for 77% of sample variation and corresponds to developmental stage, while PC2 accounts for 10% of variation and separates samples by limb type (FL vs. HL) (Figure 1E). Together, PC1 and PC2 group samples into six discrete clusters based on developmental stage and limb type.

We identified a set of 978 genes that are differentially expressed between FL and HL at HH21, HH25, and/or HH29 (“FL/HL” gene set; Table 1, Supplemental Figure 1, and Supplemental Table 1). Within the FL/HL gene set, only 12 genes are strongly enriched (defined here as an absolute value Log2FC > 2) in a FL- or HL-specific manner (Figure 1F), all of which are TFs that have been implicated in vertebrate limb development to varying degrees (Takeuchi et al., 1999, Rodriguez-Esteban et al., 1999, Szeto et al., 1999, Logan and Tabin, 1999, Zakany and Duboule, 2007, Tarchini et al., 2006, Capellini et al., 2006, Boulet and Capecchi, 2004, Wellik, 2003, Davis and Capecchi, 1994, Capellini et al., 2010, Narkis et al., 2012, Feenstra et al., 2012). In contrast, the majority (86-95%) of differentially expressed genes changed less than 2-fold (Log2FC < 1) between FL and HL (Figure 1F). These results suggest that, during early pigeon development, a small set of large gene expression changes is coupled with a large set of subtle gene expression changes to differentiate FL and HL identities. Notably, the number and level of gene expression changes that distinguish pigeon FL and HL is remarkably similar to what was recently reported for mouse FL and HL buds (Nemec et al., 2017), despite the radical differences between FL and HL morphology in pigeons and the more modest differences between limbs in mice.

**Table 1.** Summary of pigeon differential expression gene sets.

|  |  | Differential expression gene sets |  |  |
| --- | --- | --- | --- | --- |
|  |  | FL/HL | grouse | muff |
| Comparison |  | FL vs. Scaled HL | Scaled HL vs. Grouse HL | Scaled HL vs. Muff HL |
| Total | DE genes | 978 | 1425 | 3202 |
|  | Shared between grouse and muff | - | 42% (601/1425) | 19% (601/3202) |
|  | Shared with FL/HL | - | 8% (108/1425) | 10% (334/3202) |
| TFs | DE genes | 122 | 50 | 105 |
|  | Shared between grouse and muff | - | 42% (21/50) | 20% (21/105) |
|  | Shared with FL/HL | - | 18% (9/50) | 38% (40/105) |
| ECM | DE genes | 81 | 35 | 92 |
|  | Shared between grouse and muff | - | 43% (15/35) | 16% (15/92) |
|  | Shared with FL/HL | - | 26% (9/35) | 29% (27/92) |

### Dynamic *TBX5* and *PITX1* expression differences characterize feathered HL development

We previously demonstrated that in pigeons, the grouse phenotype is associated with a *cis*-regulatory mutation that reduces *PITX1* expression in the HH25 HL, while the muff phenotype is associated with *cis*-regulatory mutations that cause a combination of *PITX1* reduction and ectopic *TBX5* expression in the HH25 HL (Domyan et al., 2016, Boer et al., 2017). To determine if *PITX1* and *TBX5* expression differs in scaled, grouse, and muff HLs at other stages of development, we measured *PITX1* and *TBX5* expression in HH21, HH25, and HH29 HL buds by quantitative reverse-transcriptase PCR (qRT-PCR) (Supplemental Figure 2). We found that *TBX5* is ectopically expressed in muff HLs relative to scaled and grouse HLs at all stages analyzed. *PITX1* is significantly reduced in muff HLs relative to scaled HLs at all stages analyzed (Supplemental Figure 2). In contrast, *PITX1* is significantly reduced in grouse HLs relative to scaled HLs only at HH25 and HH29 (Supplemental Figure 2), suggesting that the temporal dynamics of *PITX1* HL expression is different between grouse and muff pigeon breeds.

### *PITX1*- and *TBX5*-dependent gene expression during feathered HL development

We predicted that the divergent *PITX1* and *TBX5* expression profiles during development of scaled, grouse, and muff HLs could reveal gene expression changes downstream of *TBX5* and/or *PITX1* that are associated with limb diversification. To test this hypothesis, we performed RNA-seq on limb buds from grouse and muff embryos isolated at HH21, HH25, and HH29 (Figure 2A). We reasoned that comparison of differentially expressed gene sets from scaled vs. grouse HLs (“grouse” gene set) and scaled vs. muff HLs (“muff” gene set) would enable us to distinguish *PITX1*-regulated gene expression changes (misexpressed in both grouse and muff HLs) from the effects of *TBX5* and/or *PITX1 + TBX5* co-regulation (misexpressed only in muff HLs).

**Figure 2.**
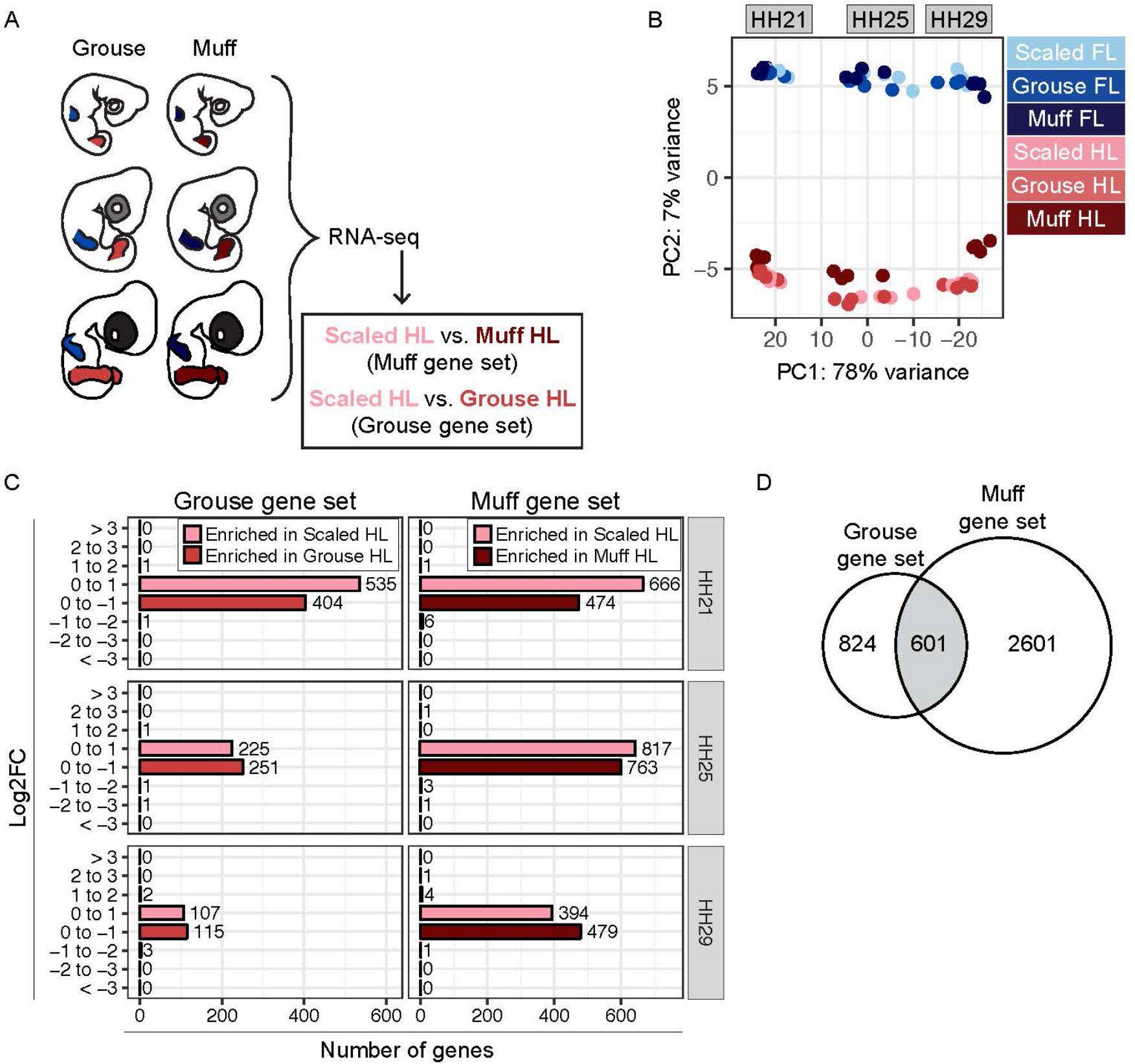
Comparison of hindlimb transcriptomes from scaled, grouse, and muff pigeon embryos. (**A**) Experimental approach to identify differentially expressed genes from grouse and muff limb buds at HH21, HH25, and HH29. (**B**) PCA based on top 500 most variable genes from all FL and HL limb bud samples. PC1 (78% of variance) separates samples by developmental stage, PC2 (7% of variance) separates samples by FL vs. HL identity. (**C**) Number of genes enriched in scaled HL or feathered HL specific manner at HH21, HH25, or HH29. Genes are grouped based on Log2FC from scaled HL vs. grouse HL or scaled HL vs. muff HL comparisons. (**D**) Venn diagram comparing differentially expressed gene sets from scaled HL vs. grouse HL and scaled HL vs. muff HL comparisons.

To test for consistency with our qRT-PCR results, we first examined the expression profiles of *PITX1* and *TBX5* in the RNA-seq samples from scaled, grouse, and muff limb buds. As expected, *TBX5* is highly expressed in all FL samples at all developmental stages with no significant difference in expression among FLs from scaled, grouse, or muff embryos (Supplemental Figure 3). Within HL samples, *TBX5* is significantly enriched in muff HLs relative to scaled and grouse HLs at all stages (Supplemental Figure 3). *PITX1*, which is expressed only in HLs, is significantly reduced in muff HLs relative to scaled HLs at all stages (Supplemental Figure 4). In contrast, *PITX1* expression is reduced in grouse HLs relative to scaled HLs only at HH25 and HH29 (Supplemental Figure 4). Thus, our analyses of *PITX1* and *TBX5* expression by qRT-PCR and RNA-seq yielded qualitatively identical results.

To determine the relationship between scaled, grouse, and muff limb bud transcriptomes, we performed PCA as described above for limb buds from scale-footed pigeons. Addition of the grouse and muff datasets had a minimal effect on the overall structure of the results; FL and HL samples remain clustered by limb type and developmental stage (Figure 2B). Within each HL cluster, muff HLs separated slightly from scaled and grouse HLs and clustered closer to their respective stage-matched FL clusters on PC2 (Figure 2B).

Next, we performed differential expression analyses to identify transcriptional changes associated with altered *PITX1* and/or *TBX5* expression in feathered HLs. Relative to scaled HLs, a total of 1425 genes are misexpressed in grouse HLs (grouse gene set) and 3202 genes are misexpressed in muff HLs (muff gene set) at HH21, HH25, and/or HH29 (Table 1, Supplemental Figure 5, and Supplemental Tables 2-3). Similar to the FL vs. scaled HL differential expression results, feathered HL development is characterized by extensive low-level gene expression changes, with nearly all differentially expressed genes (>99%) altered by less than 2-fold (Log2FC < 1, Figure 2C).

By comparing the grouse and muff gene sets, we identified 601 genes that are putatively *PITX1*-regulated, as they are misexpressed in both grouse and muff HLs (Figure 2D, Table 1, and Supplemental Table 4). A subset (96/601) of the *PITX1*-regulated genes have previously been identified as *PITX1* targets in genomic analyses of mouse *Pitx1^-/-^* HLs (Wang et al., 2018, Nemec et al., 2017), including the TFs *EMX2, HOXD11, RFX4*, *TBX15,* and *ZEB2* (Supplemental Figure 6 and Supplemental Table 4). Other *PITX1*-regulated genes encode components of several important developmental signaling pathways, as well as a variety of extracellular matrix (ECM) proteins (see below). The *PITX1*-regulated gene set also includes genes with known roles in limb development such as *short stature homeobox* (*SHOX*) (Decker et al., 2011, Tiecke et al., 2006) that, to our knowledge, were not known to be regulated by *PITX1*. Notably, *Shox* is not present in the mouse genome (but is present in other mammalian genomes, including human), precluding the identification of *Shox* as a *Pitx1* target in previous genomic analyses of mouse *Pitx1^-/-^* HLs (Wang et al., 2018, Nemec et al., 2017).

In addition to the *PITX1*-regulated genes that are misexpressed in both grouse and muff HLs, we identified 824 genes that are misexpressed only in grouse HLs (Figure 2D). This finding raises the possibility that 1) distinct genetic programs are activated depending on *PITX1* expression level (*PITX1* is differentially expressed between grouse and muff HLs; Supplemental Figure 4), or 2) additional genetic changes modify the grouse phenotype but were not detected in our whole-genome scans for allele frequency differentiation (Domyan et al., 2016).

We also identified 2601 genes that are misexpressed only in muff HLs (Figure 2D). These genes are likely regulated by *TBX5*, although we cannot rule out co-regulation by *TBX5* and *PITX1*. In summary, by comparing the transcriptional profiles associated with scaled, grouse, and muff HL development, we discovered sets of genes that are putative targets of *PITX1* and/or *TBX5* during pigeon limb development. While some of the genes we identified are known targets of *PITX1* or *TBX5*, many were not previously linked to *PITX1*, *TBX5,* or limb development.

### Partial co-option of pigeon FL genetic program during feathered HL development

We next wanted to know if feathered HL development is accompanied by FL-like transcriptional changes in genes other than *PITX1* and *TBX5*. Therefore, we probed the grouse and muff gene sets for genes that are also differentially expressed between FL and scaled HL (i.e., in the FL/HL gene set).

We first identified signatures of FL identity associated with reduced *PITX1* expression by comparing the grouse and FL/HL gene sets. Of the genes that are misexpressed in grouse HLs, ~8% (108/1425) are also differentially expressed between FL and scaled HL (Figure 3A, Table 1, and Supplemental Table 2). Within this set of 108 genes, we identified two categories of genes with FL-like gene expression patterns: FL-like Group 1 genes are enriched in FLs and grouse HLs relative to scaled HLs, while FL-like Group 2 genes have reduced expression in FLs and grouse HLs relative to scaled HLs (Figure 3B-D). FL-like Group 1 includes the TFs *HOXD11* and *LMX2*, as well as *DUSP6*, a negative regulator of FGF signaling (Figure 3B-C). FL-like Group 2 includes *PITX1*, other TFs such as *VSX1* and *RFX4*, *BMP5*, and several genes that encode cell adhesion and cytoskeletal proteins (including *LVRN*, *DSC1*, *VCAM1*, and *SPTBN5*) (Figure 3B,D).

**Figure 3.**
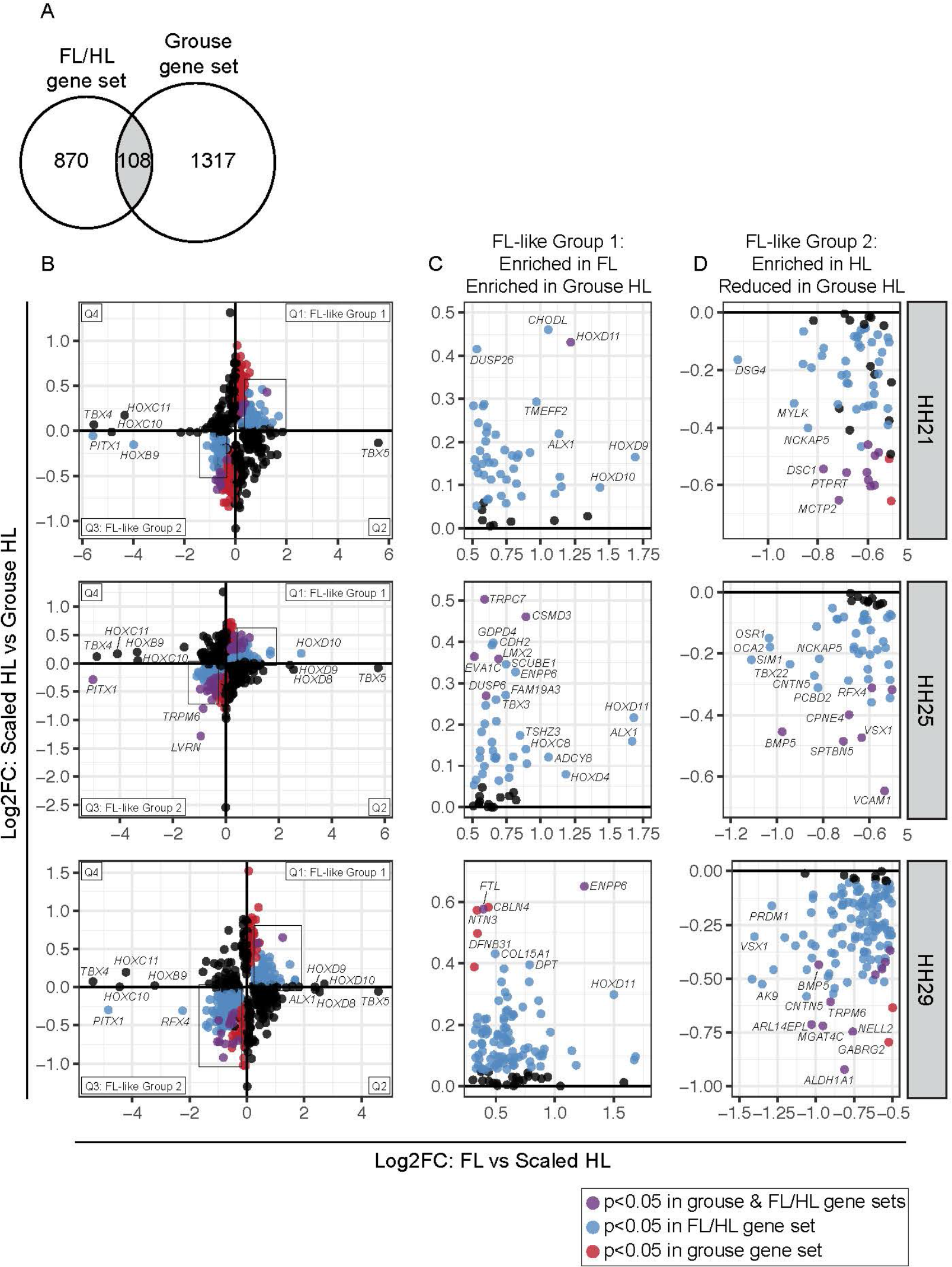
A subset of genes in the grouse HL transcriptome display FL-like expression patterns. (**A**) Venn diagram comparing differentially expressed gene sets from FL vs. scaled HL and scaled HL vs. grouse HL comparisons. (**B**) Scatterplots of Log2FC values for all genes that are differentially expressed in FL vs. scaled HL or scaled HL vs. grouse HL at HH21, HH25, or HH29. Scatterplots are divided into four quadrants (Q); Q1 and Q3 highlight genes that show FL-like expression pattern in grouse HLs. Genes in Q1 (FL-like Group 1) and Q3 (FL-like Group 2) are color coded: blue dots denote genes that are significantly differentially expressed between FL and scaled HL; red dots denote genes that are significantly differentially expressed between scaled HL and grouse HL; purple dots represent genes that are significantly differentially expressed in both comparisons. Boxes indicate regions of scatterplot displayed in (C) or (D). (**C**) Zoomed in view of FL-like Group 1 genes. (**D**) Zoomed in view of FL-like Group 2 genes. Statistical significance = adjusted p-value < 0.05.

We next identified signatures of FL identity that are putatively regulated by *TBX5* (or co-regulated by *TBX5* and *PITX1*) by comparing the muff and FL/HL gene sets. Of the genes that are misexpressed in muff HLs, ~10% (334/3202) are also differentially expressed between FL and scaled HL (Figure 4A, Table 1, and Supplemental Table 3). Similar to the grouse gene set, we identified two categories of genes with FL-like gene expression patterns (Figure 4B-D). FL-like Group 1 (enriched in FL and muff HL relative to scaled HL) includes the TFs *TBX5*, *HOXD11*, *GBX2*, *ALX1*, and *EMX2*, as well as the Wnt-associated scaffolding gene *DAAM1* (Figure 4B-C). FL-like Group 2 (reduced in FL and muff HL relative to scaled HL) includes the TFs *PITX1*, *PAX9*, *VSX1*, *RFX4*, *NKX3-2*, *DACH2* and *ZIC3*; components of developmental signaling pathways with known roles in limb development such as *WNT16*, *RDH10* and *BMP5*; and several genes encoding ECM proteins (Figure 4B,D). In addition, many other genes display non-significant FL-like expression patterns in muff HLs, including the TFs *ISL1, SOX2, HOXB5, HOXD3, HOXD4, HOXC8, HOXD8, HOXD9, HOXD10,* and *HOXD11* (Figure 4B,D).

**Figure 4.**
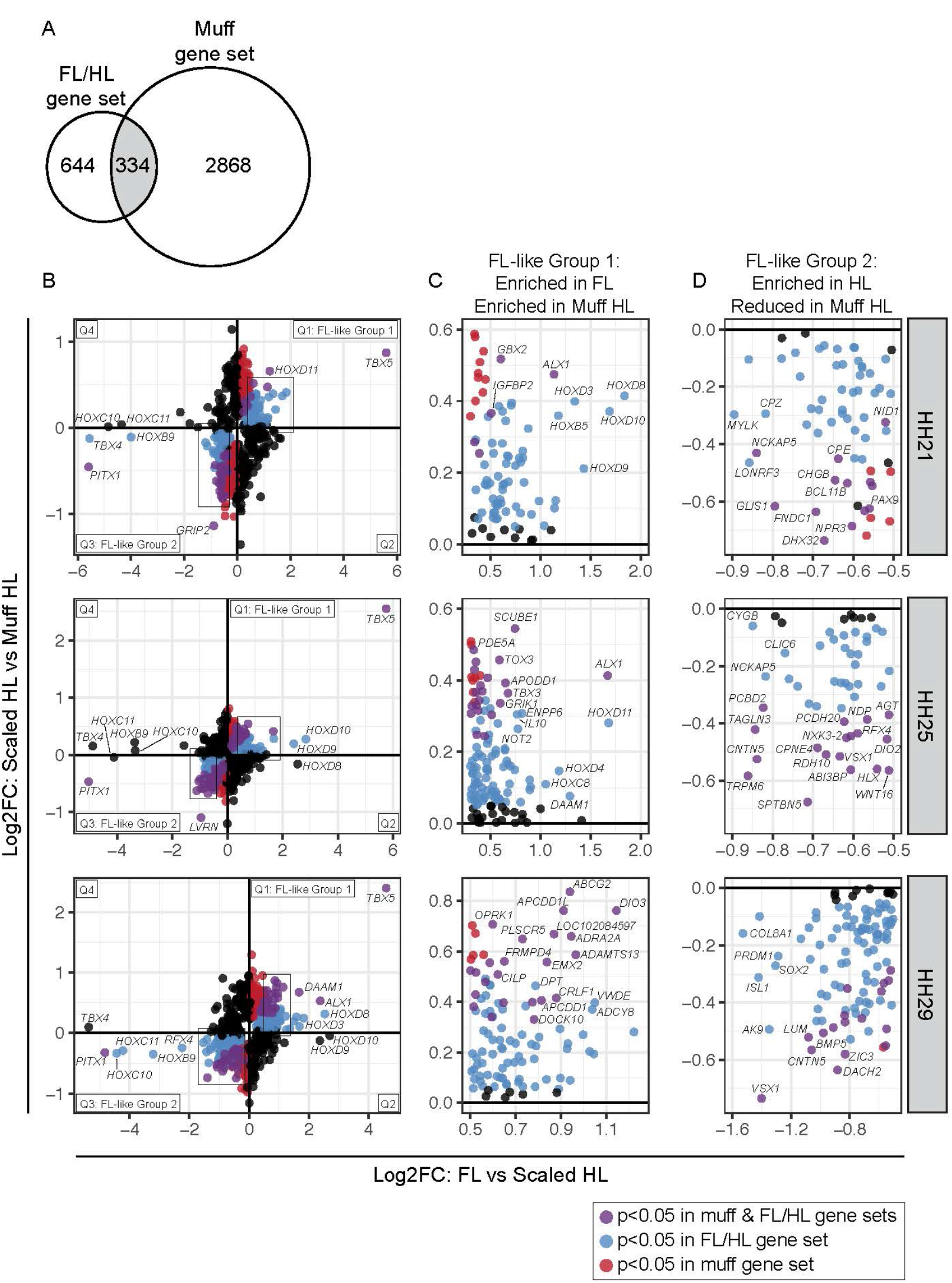
A subset of genes in the muff HL transcriptome display FL-like expression patterns. (**A**) Venn diagram comparing differentially expressed gene sets from FL vs. scaled HL and scaled HL vs. muff HL comparisons. (**B**) Scatterplots of Log2FC values for all genes that are differentially expressed in FL vs. scaled HL or scaled HL vs. muff HL at HH21, HH25, or HH29. Scatterplots are divided into four quadrants (Q); Q1 and Q3 highlight genes that show FL-like expression pattern in muff HLs. Genes in Q1 (FL-like Group 1) and Q3 (FL-like Group 2) are color coded: blue dots denote genes that are significantly differentially expressed between FL and scaled HL; red dots denote genes that are significantly differentially expressed between scaled HL and muff HL; purple dots represent genes that are significantly differentially expressed in both comparisons. Boxes indicate regions of scatterplot displayed in (C) or (D). (**C**) Zoomed in view of FL-like Group 1 genes. (**D**) Zoomed in view of FL-like Group 2 genes. Statistical significance = adjusted p-value < 0.05.

### TFs are differentially expressed in feathered HLs

In our analyses of the grouse and muff gene sets, we noted an enrichment of FL-like expression of genes encoding TFs, ECM proteins, and components of developmental signaling pathways. Hence, these gene classes could be key parts of the co-opted FL limb identity program in feathered HLs. To test this prediction, we next focused in on differentially expressed genes encoding TFs, ECM proteins, and developmental signaling components.

To identify TFs that are normally differentially expressed between FL and HL, we compared the FL/HL gene set to a genome-wide list of 1473 TFs (AnimalTFDB 2.0 database (Zhang et al., 2015)). From this list, 122 TFs are differentially expressed between FLs and scaled HLs (Figure 5A and Table 1). As expected, we detected strong FL enrichment of many known FL-specific TFs, including *TBX5, HOXD10, HOXD8, HOXD9,* and *ALX1* (Figure 5A). Likewise, we detected HL enrichment of known HL-specific TFs, including *TBX4, PITX1, HOXC11, HOXC10*, *HOXB9,* and *ISL1* (Figure 5A). Many other TFs showed FL or HL expression bias to a lesser degree, and some TFs displayed temporally dynamic FL/HL expression patterns (Figure 5A).

**Figure 5.**
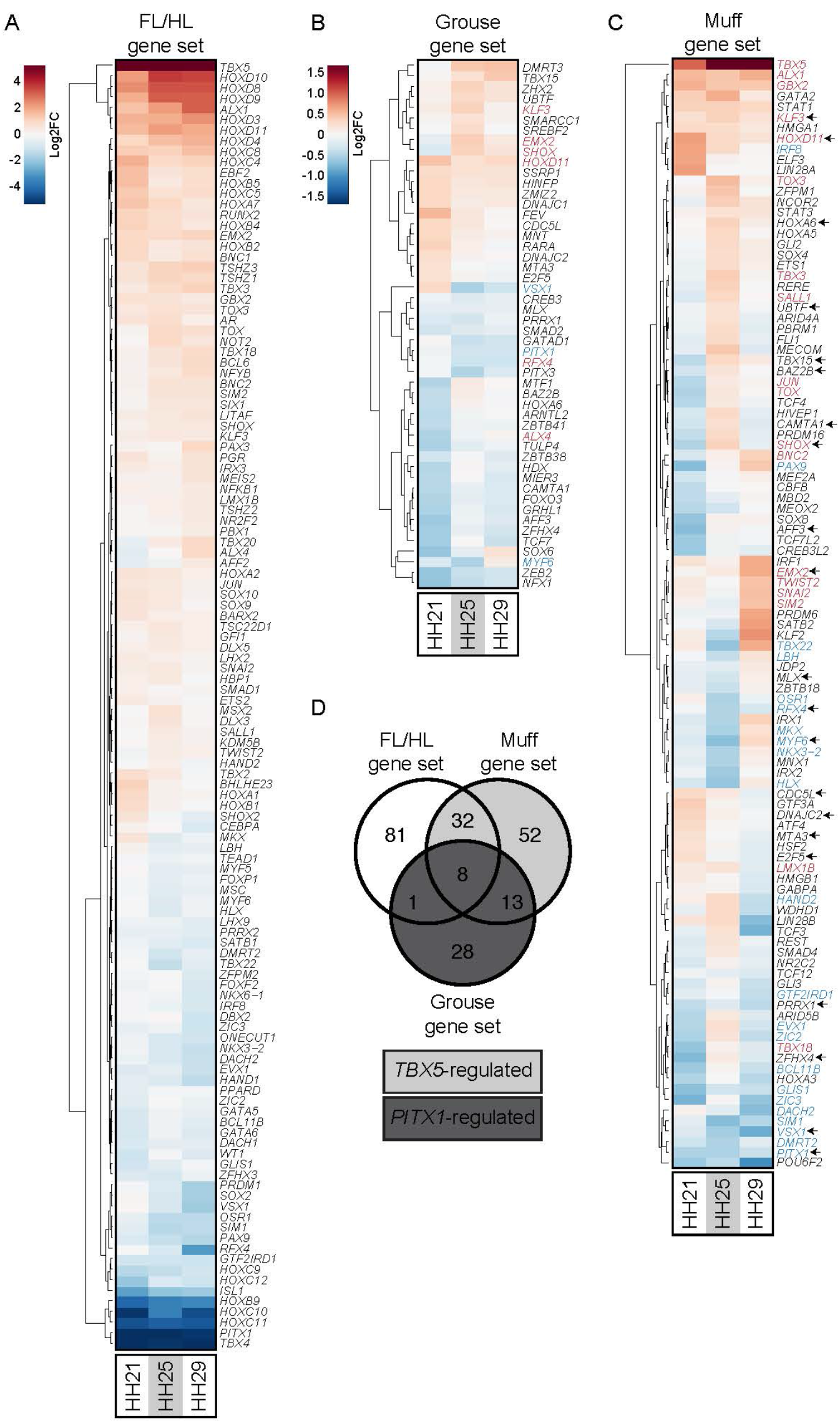
Differential expression of transcription factors associated with pigeon limb identity. (**A-C**) Gene-wise hierarchical clustering heat maps of transcription factors (TFs) that are significantly differentially expressed between FL vs. scaled HL (A), scaled HL vs. grouse HL (B), or scaled HL vs. muff HL (C). For (A-C), all TFs that are significantly differentially expressed at one or more stages are included in the heat map. The scale for (A) is Log2FC −5 (HL-enriched) to 5 (FL-enriched); the scale for (B) and (C) is Log2FC −1.5 (scaled HL-enriched) to 1.5 (grouse HL-enriched). In (B) and (C), gene names highlighted in red or blue denote genes from FL vs. scaled HL comparison that are significantly FL-enriched or HL-enriched, respectively. In (C), arrows next to gene names highlight genes that are differentially expressed in both grouse and muff HLs relative to scaled HLs. (**D**) Venn diagram showing overlap of differentially expressed TFs from FL vs. scaled HL, scaled HL vs. grouse HL, and scaled HL vs. muff HL comparisons. Dark gray shading highlights TFs that are differentially expressed in all three comparisons and are putatively regulated by *PITX1*. Light gray shading denotes TFs that are differentially expressed only in FL vs. scaled HL and scaled HL vs. muff HL comparisons, suggesting regulation by *TBX5*.

We next identified TFs that are misexpressed during feathered HL development by comparing the grouse and muff gene sets to the genome-wide TF list (Figure 5B-C). We found 50 TFs that are misexpressed in grouse HLs and 105 TFs that are misexpressed in muff HLs (Figure 5B-C and Table 1). Comparison of the grouse and muff results revealed 21 TFs that are likely regulated by *PITX1*, as they are misexpressed in both grouse and muff HLs (Figure 5C-D and Table 1). A subset of the TFs in the grouse and muff gene sets were also identified in the FL/HL gene set, suggesting that these TFs are co-opted from the normal FL identity program (Figure 5B-D and Table 1). In addition, we found a set of TFs, including several HOX and T-box genes (*HOXA3, HOXA5, HOXA6, TBX15, IRX2*), that are misexpressed in feathered HLs but are not differentially expressed between FL and scaled HLs (Figure 5B-D). These results suggest that, in addition to a partial redeployment of the FL identity program, feathered HL development involves the activation of TFs not normally required for differentiation of FLs and scaled HLs.

### Differentially expressed ECM components in feathered HLs

In birds, ECM composition differs between scaled and feathered epidermis (Sengel, 1990). Consistent with this observation, recent transcriptomic analyses have identified specific ECM genes that are differentially expressed between these epidermal types in developing archosaur embryos (Wu et al., 2018, Musser et al., 2018, Lai et al., 2018). With the importance of the ECM in mind, we identified ECM genes putatively regulated by *PITX1* and/or *TBX5* during pigeon limb development by cross-referencing our FL/HL, grouse, and muff gene sets to a list of 491 genes annotated as ECM components in the Gene Ontology database (GO:0031012) (Ashburner et al., 2000, The Gene Ontology, 2017, Carbon et al., 2009). From this list, we identified 81, 35, and 92 ECM genes in the FL/HL, grouse, and muff gene sets, respectively (Figure 6A-D and Table 1). By comparing the ECM genes in the grouse and muff gene sets, we pinpointed ECM genes that are downstream of *PITX1* and/or *TBX5* during early pigeon limb development, including ADAMTS proteases, collagens, and Wnt pathway ligands (Figure 6D). Notably, almost half (12/27) of the ECM genes shared between the FL/HL, grouse, and muff gene sets were also found to be differentially expressed between feather- and scale-forming epidermis in chick embryos (Wu et al., 2018), supporting the model that distinct ECM signatures accompany the development of divergent limb identities in birds.

**Figure 6.**
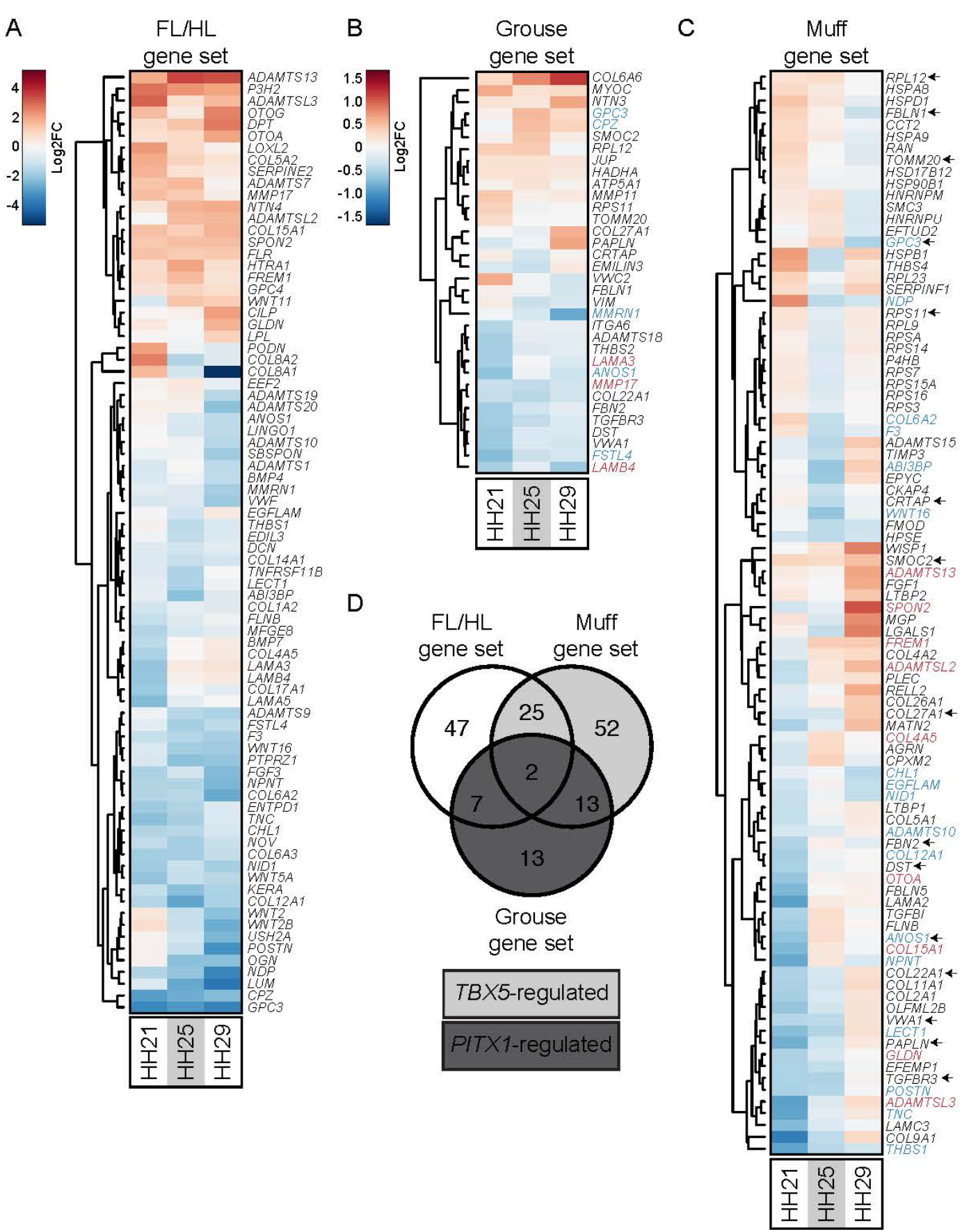
Differential expression of genes encoding extracellular matrix components is associated with pigeon limb identity. (**A-C**) Gene-wise hierarchical clustering heat maps of genes encoding extracellular matrix (ECM) components that are significantly differentially expressed between FL vs. scaled HL (A), scaled HL vs. grouse HL (B), or scaled HL vs. muff HL (C). For (A-C), all ECM-encoding genes that are significantly differentially expressed at one or more stages are included in the heat map. The scale for (A) is Log2FC −5 (HL-enriched) to 5 (FL-enriched); the scale for (B) and (C) is Log2FC −1.5 (scaled HL-enriched) to 1.5 (muff HL-enriched). In (B) and (C), gene names highlighted in red or blue denote genes from FL vs. scaled HL comparison that are significantly FL-enriched or HL-enriched, respectively. In (C), arrows next to gene names highlight genes that are differentially expressed in both grouse and muff HLs relative to scaled HLs. (**D**) Venn diagram showing overlap of differentially expressed ECM-encoding genes from FL vs. scaled HL, scaled HL vs. grouse HL, and scaled HL vs. muff HL comparisons. Dark gray shading highlights ECM-encoding genes that are differentially expressed in all three comparisons and are putatively regulated by *PITX1*. Light gray shading denotes ECM-encoding genes that are differentially expressed only in FL vs. scaled HL and scaled HL vs. muff HL comparisons, suggesting regulation by *TBX5*.

### Modulation of developmental signaling pathways in feathered HLs

Normal limb development requires the integration of several key developmental signaling pathways including the SHH, FGF, BMP, RA, and Wnt pathways (reviewed in (Duboc and Logan, 2011b, Tickle, 2015)). Within the FL/HL gene set, we identified suites of genes involved in each of these pathways using Ingenuity Pathway Analysis (IPA) (Supplemental Figure 7), suggesting that fine-tuning of developmental pathways is a normal part of FL vs. HL development. To determine if misexpression of *PITX1* and/or *TBX5* in feathered HLs is associated with altered expression of genes within the same pathways, we also probed the grouse and muff gene sets using IPA. In both the grouse and muff gene sets, we detected signatures of differential FGF, RA, and Wnt signaling (Supplemental Figure 7), providing evidence that *PITX1* interacts with each of these pathways. The muff gene set also includes signatures of differential BMP and SHH signaling (Supplemental Figure 7), suggesting that *TBX5*, and not *PITX1*, is a major regulator of BMP and SHH signaling during early stages of limb development (Supplemental Figure 7). Together, these analyses suggest that fine-tuning of developmental signaling pathways by *PITX1* and *TBX5* is associated with the development of distinct limb morphologies.

### Evidence for conservation of limb identity programs among birds

Despite highly variable adult limb morphologies, the genetic toolkit that regulates limb development among vertebrates is evolutionarily conserved. However, the degree to which regulation of these genes is conserved across distantly related species remains unclear. To determine if limb-specific transcriptional profiles are conserved across large phylogenetic distances, we first compared gene expression in limb buds from pigeons and chickens, two distantly related avian species. We generated an RNA-seq dataset from FL and HL buds from scale-footed chicken embryos at developmental stages comparable to our pigeon dataset (HH21, HH25, and HH29). Consistent with the pigeon dataset, PCA based on the 500 most differentially expressed genes separated samples by stage (PC1, 82% of variance explained) and limb type (PC2, 11%) (Figure 7A). We identified a set of 2160 genes that are differentially expressed between chicken FL and HL buds at HH21, HH25, and/or HH29 (chicken FL/HL gene set, Supplemental Figure 8 and Supplemental Table 5). As in our pigeon analyses, we found that most of the gene expression changes between chicken FL and HL are relatively modest (Log2FC < 1) (Figure 7B). Similar to our analysis of pigeon FL and HL buds, we identified a suite of TFs, genes encoding ECM proteins, and components of developmental signaling pathways that are differentially expressed between chicken FL and HL buds (Supplemental Figure 9).

**Figure 7.**
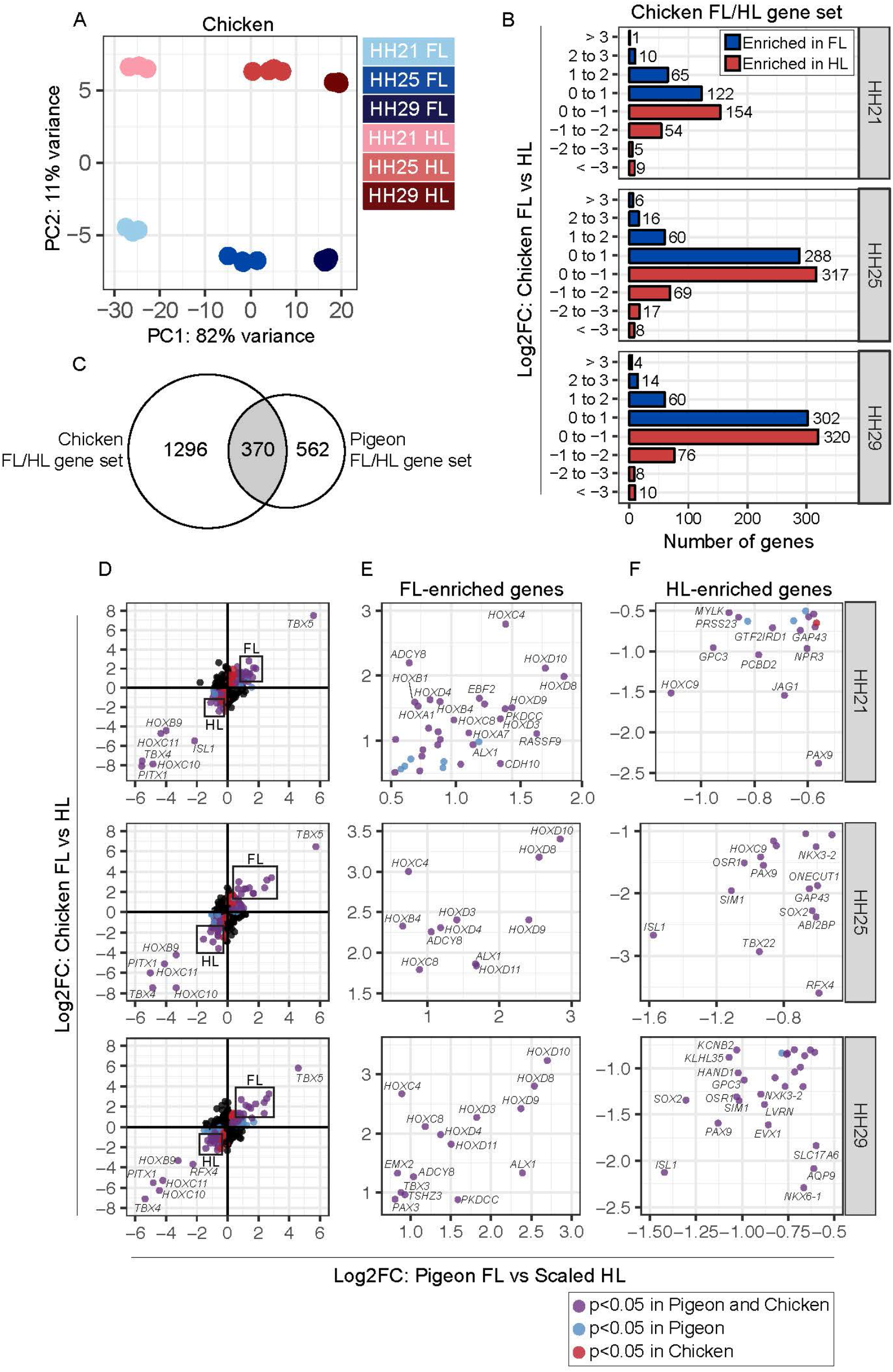
Identification of conserved gene expression patterns in embryonic limb buds from chicken and pigeon. (**A**) PCA based on top 500 most variable genes in chicken FL and scaled HL buds at HH21, HH25, and HH29. PC1 (82% of variance) separates samples by stage; PC2 (11% of variance) separates samples by limb type. (**B**) Number of genes that show FL-enriched or scaled HL-enriched expression pattern in chicken embryos at HH21, HH25, or HH29. Genes are grouped based on Log2FC. (**C**) Venn diagram showing overlap of chicken FL vs. HL and pigeon FL vs. scaled HL gene sets. (**D**) Scatterplots of Log2FC values for all genes that are differentially expressed in chicken FL vs. HL or pigeon FL vs. scaled HL at HH21, HH25, or HH29. Scatterplots are divided into four quadrants (Q); Q1 and Q3 highlight genes that show FL-enriched or HL-enriched expression in both avian species. Genes in Q1 (FL-enriched) and Q3 (HL-enriched) are color coded: blue dots denote genes that are significantly differentially expressed in pigeon FL vs. scaled HL; red dots denote genes that are significantly differentially expressed in chicken FL vs. HL; purple dots represent genes that are significantly differentially expressed in both comparisons. Boxes indicate regions of scatterplot displayed in (C) or (D). (**C**) Zoomed in view of FL-enriched genes. (**D**) Zoomed in view of HL-enriched genes. Statistical significance = adjusted p-value < 0.05.

To determine the degree to which the limb identity network is conserved between pigeons and chickens, we compared the FL/HL gene sets from each species. For this analysis, we included only the genes that are annotated in both species (based on Cliv_2.1 (Holt et al., 2018) and Gallus_gallus-5.0 (Warren et al., 2017)), resulting in comparison of 933/978 pigeon and 1667/2160 chicken genes. Despite this limitation, we found 370 genes that are differentially expressed between FL and HL in both chicken and pigeon (Figure 7C and Supplemental Table 6). In both species, *TBX5* stood out as the most FL-enriched gene, while TFs *HOXD8, HOXD9, HOXD10*, and *ALX1* were also highly FL-enriched (Log2FC > 2) in both species (Figure 7D-E). A small set of TFs (*TBX4, PITX1, HOXB9, HOXC10, HOXC11, ISL1*) was highly HL-enriched (Log2FC > 2) in both species (Figure 7D,F). In both chickens and pigeons, many other TFs were expressed in a FL- or HL-specific manner to a lesser degree (Figure 7D-F and Supplemental Table 6). Notably, 20% (76/370) of the genes identified in both the pigeon and chicken FL vs. HL datasets were TFs (Supplemental Table 6), further suggesting that TFs are highly conserved components of limb identity programs.

### Evidence for conservation of limb identity programs among amniotes

Recent studies have characterized embryonic FL and HL bud transcriptomes in several mammalian species, including mouse, opossum, pig, and bat (Taher et al., 2011, Maier et al., 2017, Sears et al., 2018, Wang et al., 2018, Gyurjan et al., 2011, Eckalbar et al., 2016, Shou et al., 2005, Nemec et al., 2017). However, similar analyses in other vertebrate classes are sparse, precluding broader comparative analyses of limb identity networks. We therefore extended our analyses by identifying differentially expressed genes in embryonic FLs and HLs of brown anole lizards (*Anolis sagrei*, Supplemental Figure 10). In addition, we re-analyzed several publicly available mammalian (mouse and opossum) limb bud datasets (Nemec et al., 2017, Maier et al., 2017, Amandio et al., 2016). Together with our pigeon and chicken datasets, these additional data enabled us to identify conserved components of the limb identity network in Aves (pigeon and chicken), Sauropsida (pigeon, chicken, and anole), Mammalia (mouse and opossum), and Amniota (pigeon, chicken, anole, mouse, and opossum) (Figure 8A). To facilitate comparisons among species, we categorized samples from each dataset into ridge, bud, or paddle developmental stages, which approximately correspond to chicken stages HH21, HH25, and HH29, respectively.

**Figure 8.**
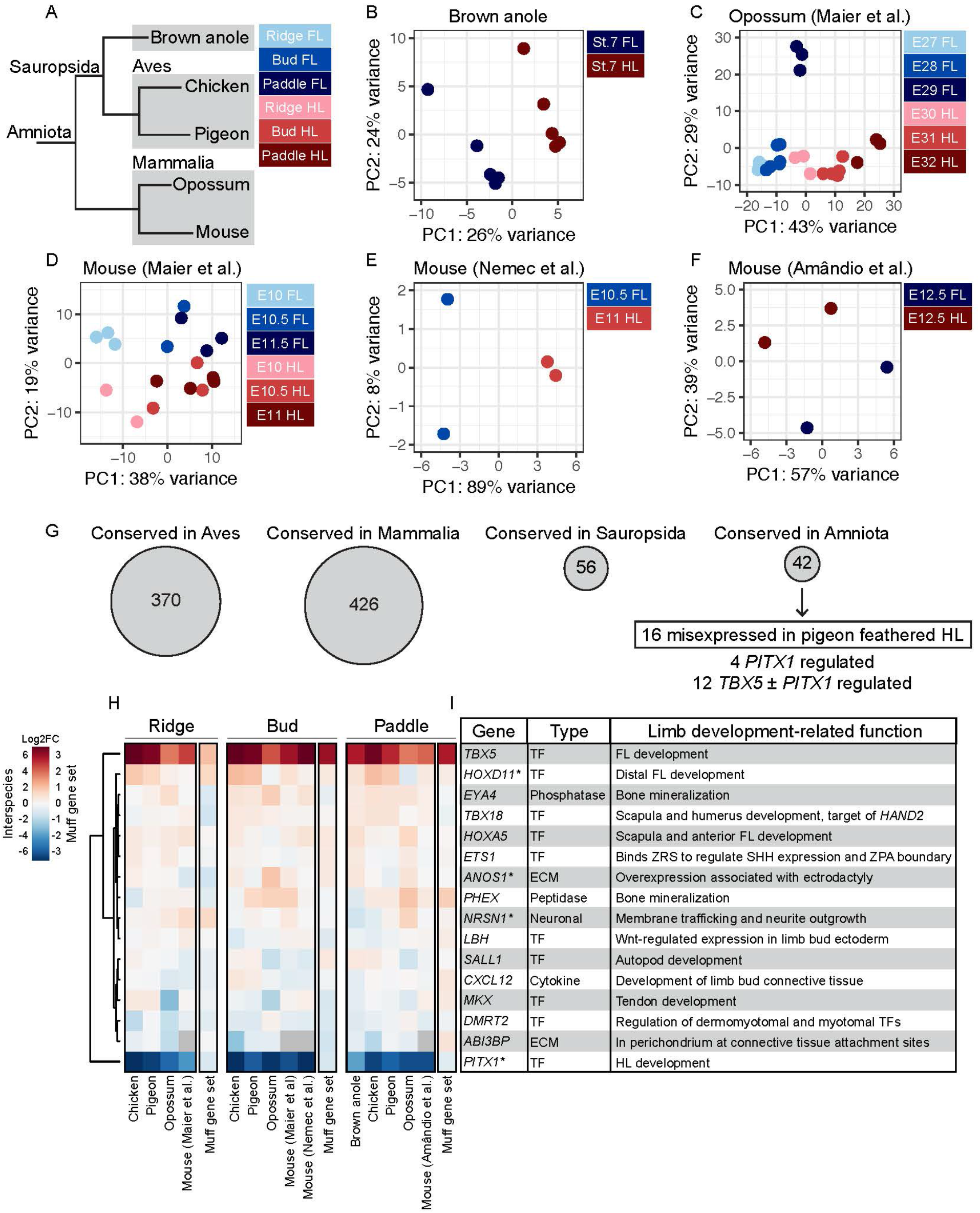
Identification of deeply conserved elements of limb identity program in amniotes. (**A**) Tree summarizing evolutionary relationships between species included in comparative transcriptomics analyses. To facilitate analyses, samples from all datasets were categorized into three stage groups: ridge, bud, or paddle. (**B-F**) PCAs based on top 500 most variable genes from brown anole (B), opossum (C), and three different mouse (D-F) limb bud datasets. (C) and (D) are analyses of datasets from (Maier et al., 2017), (E) is analysis of dataset from (Nemec et al., 2017), and (F) is analysis of dataset from (Amandio et al., 2016). For (B-F), specific stages of samples are listed and color coded based on stage group. (**G**) Summary of number of genes differentially expressed in FL vs. HL of Aves (chicken and pigeon), Mammalia (mouse and opossum), Sauropsida (brown anole and chicken or pigeon), and Amniota (brown anole, chicken or pigeon, and mouse or opossum). Of the 42 genes differentially expressed in FL vs. HL across amniotes, 16 are also misexpressed in pigeon feathered HLs. Of these, 4 genes are putative *PITX1* targets (misexpressed in grouse and muff HLs relative to scaled HLs) and 12 are likely regulated by *TBX5* or co-regulated by *TBX5* and *PITX1* (misexpressed in muff HLs, but not grouse HLs). (**H**) Gene-wise hierarchical clustering heat map of 16 genes that are differentially expressed in FL vs. HL across amniotes and scaled vs. feathered HL in pigeons. The scale for between-species comparisons is Log2FC −6 (HL-enriched) to 6 (FL-enriched); the scale for pigeon HL comparisons is Log2FC −3 (scaled HL-enriched) to 3 (muff HL-enriched). Gray boxes within the heat map denote no data (*ABI3BP* is not annotated in mouse genome). (**I**) Table summarizing 16 genes included in (**H**), including type of gene, limb development-related function (if known) with corresponding references. Asterisks (*) denote 4 putative *PITX1* targets.

Similar to the avian PCA results, limb buds from anole, mouse, and opossum datasets generally clustered based on limb type (FL vs. HL) and stage (for datasets that included multiple stages) (Figure 8B-F). For each dataset, we identified genes that are differentially expressed between stage-matched FL and HL buds (Supplemental Table 7). Consistent with our results from pigeon and chicken, in most data sets, almost all gene expression changes between FL and HL are low-level (Log2FC < 1), with only a small set of highly differentially expressed genes (Log2FC > 2) (Supplemental Figure 11).

To identify deeply conserved limb identity genes, we compared the differentially expressed gene sets from pigeon, chicken, anole, mouse, and opossum. Despite caveats associated with this type of meta-analysis that undoubtedly reduce our power to detect conserved genes (addressed in Discussion), we identified differentially expressed genes that are associated with FL/HL limb identity in Mammalia (426 in mouse and opossum), Sauropsidia (56 in anole and pigeon or chicken), and Amniota (42 in anole, pigeon or chicken, and mouse or opossum) (Figure 8G and Supplemental Table 7). Within both the Sauropsidia and Amniota gene sets, ~50% (25/56 in Sauropsidia, 21/42 in Amniota) of the conserved differentially expressed genes are TFs (Supplemental Table 7), again highlighting that TFs are a highly conserved component of limb identity networks.

Finally, we asked if the genetic program associated with feathered HL development in pigeons represents a co-option of deeply conserved components of the FL identity program. We compared the grouse and muff gene sets to the 42 limb identity genes conserved in Amniota and found that 38% (16/42) of deeply conserved limb identity genes are also differentially expressed in grouse and/or muff pigeon HLs (Figure 8G-I). In addition to *PITX1*, we identified 3 deeply conserved limb identity genes (*HOXD11*, *ANOS1*, *NRSN1*) that are misexpressed in both grouse and muff HLs, suggesting that these genes are conserved downstream targets of *PITX1* (Figure 8H-I). In addition to *TBX5*, 11 deeply conserved limb identity genes *(EYA4, TBX18, HOXA5, ETS1, PHEX, LBH, SALL1, CXCL12, MKX, DMRT2, ABI3BP)* are misexpressed only in muff HLs (Figure 8H-I). This set of 11 genes, which includes 7 TFs, is putatively regulated by *TBX5* or co-regulated by *TBX5* and *PITX1*. In particular, *CXCL12*, *MKX*, and *DMRT2* may be co-regulated by *TBX5* and *PITX1*, as these genes were recently identified as conserved regulatory targets of *PITX1* in mouse and anole HLs (Wang et al., 2018). Of the 16 deeply conserved limb identity genes that are deployed during pigeon feathered HL development, most have been implicated in specific aspects of limb development with varying degrees of supporting evidence (*TBX5*, *HOXD11*, *TBX18*, *HOXA5*, *ETS1*, *LBH*, *SALL1*, *CXCL12*, *MKX*, *PITX1*) (Boulet and Capecchi, 2004, Zakany and Duboule, 2007, Sheeba and Logan, 2017, Haraguchi et al., 2015, Xu et al., 2013, Lettice et al., 2012, Kvon et al., 2016, Briegel and Joyner, 2001, Kawakami et al., 2009, Farrell and Munsterberg, 2000, Nassari et al., 2017, Odemis et al., 2005, Liu et al., 2010, Ito et al., 2010, Duboc and Logan, 2011b) (Figure 8I). For others (*EYA4*, *ANOS1*, *PHEX*, *NRSN1*, *DMRT2*, *ABI3BP*), a link to limb development has not been established. Furthermore, for most of the 16 deeply conserved limb identity genes, the precise regulatory connections to *TBX5* and/or *PITX1* remain open for discovery.

## Discussion

We identified transcriptional changes associated with differences in FL, HL, and limbs with mixed identities. We took advantage of regulatory variants associated with pigeon foot feathering to specifically determine how shifts in the expression of two regulators of limb identity, *PITX1* and *TBX5*, affect transcription of downstream genes during embryonic limb development. We found that most gene expression differences between FLs and HLs are subtle, and that only a small set of genes is highly differentially expressed. We identified suites of genes encoding TFs, ECM proteins, and components of developmental signaling pathways with known roles in limb development that are regulated downstream of *TBX5* and/or *PITX1*. Finally, by comparing limb identity networks across amniotes, we found a small group of deeply conserved gene expression changes associated with limb identity. Some of the genes in this core group are also misexpressed in feathered pigeon HLs, consistent with a partial HL to FL transformation at both the phenotypic and transcriptomic level.

### Embryonic limb identity networks

In birds, we found that limb type-specific expression of a small set of TFs is coupled with an accumulation of widespread low-level gene expression changes to form radically divergent FL and HL morphologies. Notably, transcriptomic studies of mouse embryonic limb development identified a similar number and level (i.e. fold-change) of gene expression changes between FL and HL (Nemec et al., 2017). These parallel findings are particularly interesting considering that variation in adult limb morphologies of mice (FL and HL are covered with hair and have similar skeletal proportions) is less dramatic than in birds (FLs are feather covered wings, HLs are scaled covered legs, and limb skeletons are highly divergent). Thus, the identity of the differentially expressed genes, and not the overall number or level of gene expression changes, ultimately determines adult limb phenotype. In our most phylogenetically inclusive analyses, we identified a core set of gene expression changes associated with amniote FL/HL limb identity, but we also found many genes that are differentially expressed between FLs and HLs in a species-specific manner. We predict that these unique sets of genes are associated with the development of species-specific limb phenotypes. In support of this idea, specific sets of genes initiate distinct developmental programs to form feathers or different scale types in archosaur embryos (Musser et al., 2018, Wu et al., 2018, Lai et al., 2018).

### Parsing *TBX5* vs. *PITX1*-dependent regulation of the limb identity network

Pigeon foot feathering offers a unique opportunity to understand the independent and combinatorial effects of *PITX* and *TBX5* expression changes on limb patterning. Both grouse and muff HLs are feathered, but the combination of reduced *PITX1* and ectopic *TBX5* expression (muff) had a more pronounced effect on the limb regulatory network than reduced *PITX1* expression alone (grouse). By comparing grouse and muff HL transcriptomes, and by cross-referencing these data to HL transcriptomes from *Pitx1* mouse mutants (Nemec et al., 2017, Wang et al., 2018), we were able to infer *PITX1*-specific transcriptional targets in pigeons. However, we were unable to separate *TBX5*-specific from *TBX5/PITX1*-synergistic effects in our pigeon analyses. This shortcoming could be resolved by analyzing HL bud transcriptomes from feather-footed pigeons that carry the *Slipper* (*TBX5*) allele but not the *grouse* (*PITX1*) allele, although such breeds are rare (English and Pigmy Pouters only).

### Partial co-option of pigeon FL identity program in feathered HLs

By comparing the differentially expressed genes from scaled and feather-footed pigeons, we found that the gene expression programs controlling feathered HL development are partially co-opted from the normal FL program. Thus, in feather-footed pigeons, a portion of the FL program is redeployed in the HL to form feathered HLs with partial anatomical and molecular FL identity. Spatiotemporal modulation of gene expression, particularly of genes like *PITX1* and *TBX5* that function at the top of gene regulatory hierarchies, is an important driver of evolutionary diversification (Davidson, 2006, Erwin and Davidson, 2009, Rebeiz et al., 2015, Infante et al., 2018). Generation of phenotypic diversity and novelty via co-option of genetic circuits or top-level network regulators is a repeated theme in evolution; notable examples include co-option of the adult skeletogenic network to form the larval skeleton in echinoderms, redeployment of the highly conserved arthropod appendage-patterning network to form beetle horns, and overlap of the genetic networks that control embryonic limb and external genital development in tetrapods (Rebeiz et al., 2015, Davidson, 2006, Erwin and Davidson, 2009, Infante et al., 2015, Infante et al., 2018, Gao and Davidson, 2008, Tschopp et al., 2014, Herrera and Cohn, 2014, Leal and Cohn, 2016, Leal and Cohn, 2018). Our comparative analyses in pigeons indicate that TFs are a major component of the FL identity program redeployed in feathered HLs; similarly, others have noted that genes at the top of regulatory networks are more evolutionarily stable than differentiation genes at the network periphery (Davidson, 2006, Erwin and Davidson, 2009, Rebeiz et al., 2015). Within the co-opted FL program in feathered HLs, we also identified genes encoding developmental signaling pathway components and ECM proteins, which link subcircuits of the regulatory network and function as differentiation genes at the network periphery, respectively (Erwin and Davidson, 2009, Davidson, 2006, Rebeiz et al., 2015). These findings suggest that changes in the expression of top-level regulators (*PITX1* and/or *TBX5*) have widespread effects on all levels of the limb development network.

Despite a partial co-option of the FL identity program, it is important to note that pigeon feathered HLs are not FLs. They are not the consequence of a complete redeployment of the FL identity network onto a blank canvas. Instead, the ectopic FL program is superimposed onto the resident HL regulatory network, resulting in a HL with mixed identity. During normal limb bud development, ectodermal identity is determined in response to signals from the underlying mesoderm (Cairns and Saunders, 1954, Saunders et al., 1959, Sengel, 1990, Hughes et al., 2011). In muff HLs, ectopic and spatially-restricted mesodermal expression of *TBX5* signals to the ectoderm to form feathered epidermis. The largest flight-like feathers of muffed pigeon feet develop superficial to the ectopic *TBX5* expression domain in the posterior-dorsal limb bud. This spatially-restricted FL-like genetic program occurs within the context of HL identity program with reduced *PITX1* expression. Together, changes in *PITX1* and *TBX5* expression cause a reorganization of the downstream limb identity network that results in a limb that is neither completely HL nor completely FL, as evident from the changes in both epidermal appendage type and musculoskeletal patterning (Domyan et al., 2016). This layering of multiple genetic networks represents an intriguing potential mechanism of phenotypic diversification.

### Deeply conserved components of limb identity networks

By comparing FL and HL bud transcriptomes from several amniote species, we identified a core set of deeply conserved limb identity genes that is enriched for genes encoding TFs, ECM proteins, and signaling pathway components. We suspect that only a subset of the deeply conserved genes was identified in our analyses, due primarily to technical caveats associated with cross-species genomic analyses (e.g., differences in sample preparation, sequencing, quality of genome assembly and annotation). Nevertheless, our analyses suggest that the number of genes differentially expressed between FL and HL in a species-specific manner is considerably greater than the number of deeply conserved limb identity genes. This result suggests that relatively few genes are absolutely required for amniote FL vs. HL identity and raises the possibility that there is considerable evolutionary variation in the genes that were recruited to the limb development and identity programs. Another classic example of this paradigm is animal sex determination, in which a small set of master sex determination genes are highly conserved across species (e.g. *DMRT1*, *SOX3*), but the genetic players vary dramatically at the network periphery (Herpin and Schartl, 2015).

Of the deeply conserved FL/HL identity genes we identified, a subset was also differentially expressed between scaled and feathered pigeon HLs, suggesting that the genetic program underlying pigeon foot feathering is not completely novel, but instead incorporates elements of the deeply conserved FL/HL limb identity network. In the future, it will be interesting to determine if similar mechanisms of network co-option and/or overlaying of multiple regulatory networks are broadly associated with vertebrate limb diversification.

## Materials and methods

### Animal husbandry, isolation of embryonic tissue and sample sex determination

*Columba livia* were housed in accordance with the University of Utah Institutional Animal Care and Use Committees of University of Utah (16-03010). Pigeon eggs were collected from Racing Homer (scaled), Oriental Frill (grouse) and English Trumpeter (muff) breeding pairs and incubated to the desired embryonic stage. White Leghorn chicken eggs were obtained from AA Lab Eggs (Westminster, California) and incubated to the desired stage. For avian samples, pairs of forelimb (FL) and hindlimb (HL) buds were dissected from scaled, grouse and muff pigeon embryos or chicken embryos at HH21 (embryonic day E3.5), HH25 (E4.5), and HH29 (E6) and stored in RNAlater (ThermoFisher Scientific) at −80°C. Additional tissue was harvested from each embryo and stored at −80°C for DNA extraction and genotyping. Genomic DNA was isolated from each embryonic tissue sample using a DNeasy Blood and Tissue Kit (Qiagen). For each pigeon sample, sex was determined using a previously published PCR-based assay (Fridolfsson and Ellegren 1999).

*Anolis sagrei* were maintained at the University of Georgia following published guidelines (Sanger et al., 2008a). Animals were wild individuals captured at the Fairchild Tropical Botanic Gardens in Miami, FL. Breeding cages housed up to 4 adult females and 1 adult male together. Eggs were collected from nest boxes weekly and incubated at 26°C. Pairs of FL and HL buds were dissected from Stage 7 *A. sagrei* embryos and stored in RNAlater at −80°C. Embryos were staged according to (Sanger et al., 2008b). All experiments followed the National Research Council’s Guide for the Care and Use of Laboratory Animals and were performed with the approval and oversight of the University of Georgia Institutional Animal Care and Use Committee (A2015 02-020-Y3-A13).

### RNA isolation, cDNA synthesis, and qRT-PCR analysis

Total RNA was extracted from FL and HL bud samples using the RNeasy Mini Kit with RNase-Free DNAse Set and a TissueLyser LT (Qiagen). For each sample, a cDNA library was prepared using M-MLV Reverse Transcriptase with oligo(dT) primer (ThermoFisher Scientific). Intron-spanning amplicons from pigeon *PITX1*, *TBX5*, and *ACTB* were amplified from cDNA libraries using a CFX96 instrument and iTaq Universal Sybr Green Supermix (BioRad). Primer sequences were published previously (Domyan et al., 2016). For a single qRT-PCR experiment, three technical replicates were performed for each sample and the mean value was determined. Each experiment was repeated three times. Statistical significance was determined using a pairwise Wilcoxon rank sum test in R version 3.3.3 (R Core Team, 2018).

### Library preparation and RNA sequencing of avian samples

RNA-sequencing libraries were prepared and sequenced by the High-Throughput Genomics and Bioinformatic Analysis Shared Resource at the University of Utah. RNA sample quality was assessed using the RNA ScreenTape Assay (Agilent) and a sequencing library was prepared using the TruSeq Stranded mRNA Sample Prep Kit with oligo(dT) selection (Illumina). 125-cycle paired-end sequencing was performed on an Illumina HiSeq 2500 instrument (12 libraries/lane). An average of 21 million reads were generated for each pigeon sample and 25 million reads for each chicken sample.

### Analysis of avian RNA-seq data

Sequencing read quality was assessed with FastQC (Babraham Bioinformatics). Read alignment was performed using STAR version 2.5.0a (Dobin et al., 2013). Illumina adapters were trimmed and reads were aligned to the pigeon Cliv_2.1 reference assembly (Holt et al., 2018) or chicken Gallus_gallus-5.0 genome assembly (Warren et al., 2017) with STAR using the 2-pass mode. GTF annotation files were used to guide spliced read alignments. The average percentage of uniquely mapped reads was 84.07% for pigeon samples and 91.45% for chicken samples. Mapped reads were assigned to genes using featureCounts from the Subread package version 1.5.1 (Liao et al., 2014), with an average of 72.36% of reads assigned in pigeon samples and 74.48% in chicken samples. Principal component and differential expression analyses were performed with the R package DESeq2 version 1.12.4 (Love et al., 2014). Read count files were imported into DESeq2 and a DESeqDataSet object was generated that included sample sex and sequencing lane as covariates. Prior to running any DESeq2 functions, datasets were pre-filtered to remove genes with a total of 0 reads across all samples. To generate differentially expressed gene lists, we employed a Benjamini & Hochberg adjusted p-value cutoff of p<0.05 because in some cases, more conservative cutoffs resulted in gene sets that did not include *PITX1* or other known limb identity factors in the FL vs. scaled HL comparisons. In order to perform downstream comparative analyses, differentially expressed gene lists were annotated with human gene names using a custom R script.

### Analysis of transcription factors, extracellular matrix components, and signaling pathways

To identify transcription factors (TFs) within our differentially expressed gene lists, we cross-referenced gene lists to a list of ~1400 genome-wide TFs compiled from AnimalTFDB 2.0 (Zhang et al., 2015). Extracellular matrix genes were identified by comparing differential expression results to a list of genes associated with the GO term “extracellular matrix” (GO:0031012). Ingenuity Pathway Analysis 2.2 (IPA, Qiagen) was used to identify differentially expressed components of specific developmental signaling pathways with roles in limb development. We specifically searched for significant enrichment of the following canonical pathways in IPA: Wnt signaling (Wnt/β-catenin signaling, PCP pathway, Wnt/Ca+ signaling), BMP signaling, TGF-β signaling, Sonic Hedgehog signaling, and RAR activation.

### RNA-seq and analysis of Brown anole (*Anolis sagrei*) limb buds

*A. sagrei* FL and HL RNA-seq libraries were prepared using the TruSeq Stranded mRNA Sample Prep Kit with oligo(dT) selection (Illumina). Five separate FL and HL library sets were prepared from limb RNA collected from five different embryos. 75-cycle paired-end sequencing was performed on an Illumina NextSeq 500 instrument (10 libraries/lane). An average of 43 million reads were generated for each *A. sagrei* library. Read quality assessment, adapter trimming, and mapping to the Green anole (*Anolis carolinensis*) AnoCar2.0 genome assembly were performed as described above with modified alignment parameters (--outFilterScoreMinOverLread 0.3 -- outFilterMatchNminOverLread 0.3 --outFilterMultimapNmax 15) to allow for mismatches due to species-specific sequence differences. The average percentage of uniquely mapped reads was 79.19% for Brown anole samples. Mapped reads were assigned as described above, with an average of 61.23% of reads assigned. Principal component and FL vs. HL differential expression analyses were performed using DESeq2 as described above.

### Analysis of publicly available mammalian RNA-seq datasets and comparative transcriptomics between species

Raw RNA-seq reads from three limb bud datasets from mouse (*Mus musculus*, (Maier et al., 2017, Nemec et al., 2017, Amandio et al., 2016)) and one limb bud dataset from opossum (*Monodelphis domestica*, (Maier et al., 2017)) were downloaded from the GEO database (GSE71390, GSE79028, GSE100734). Read quality assessment, adapter trimming, and read mapping to either the mouse GRCm38.p6 or opossum monDom5 genome assemblies were performed as described above, with an average alignment percentage of 86.90% (mouse, (Maier et al., 2017)), 76.46% (mouse, (Nemec et al., 2017)), 88.43% (mouse, (Amandio et al., 2016)), and 84.76% (opossum, (Maier et al., 2017)). Aligned reads were assigned to genes as described above, with an average of 66.47% (mouse, (Maier et al., 2017)), 48.80% (mouse, (Nemec et al., 2017)), 70.90% (mouse, (Amandio et al., 2016)), and 58.93% (opossum, (Maier et al., 2017)) successfully assigned reads. For each dataset, principal component and FL vs. HL differential expression analyses were performed using DESeq2 as described above. A master list of differentially expressed genes in mouse limb buds was generated by combining the gene sets obtained from the mouse datasets.

To facilitate comparative transcriptomic analyses, gene lists from all species were annotated with human gene names using a custom R script. To identify differentially expressed genes conserved in Aves or Mammalia, we intersected FL vs. HL gene lists from either pigeon and chicken (Aves) or mouse and opossum (Mammalia). FL vs. HL genes conserved in Sauropsida were determined by intersecting gene lists from pigeon, chicken, and Brown anole and identifying genes that are differentially expressed in Brown anole and at least one member of Aves (pigeon and/or chicken). Similarly, FL vs. HL genes conserved in Amniota were determined by identifying genes that are differentially expressed in Brown anole, at least one member of Aves (pigeon and/or chicken) and at least one member of Mammalia (mouse and/or opossum). Finally, we cross-referenced the Amniota gene list to the pigeon scaled HL vs. grouse HL and scaled HL vs. muff HL gene lists to identify deeply conserved FL vs. HL genes that are also misexpressed during grouse and/or muff HL development.

## Acknowledgements

We thank Robert Greenhalgh and Timothy Mosbruger for their help with genomic analyses; Timothy Parnell and the Huntsman Cancer Institute Bioinformatic Analysis Shared Resource for assistance with and access to Ingenuity Pathway Analysis; and Anna Vickrey and Robert Greenhalgh for comments on the manuscript. We acknowledge the laboratory of Mark Yandell and the Center for High Performance Computing at the University of Utah for support and computing resources.

## Data availability

All RNA-seq data generated for this study have been deposited at Gene Expression Omnibus under accession numbers GSE127775 (*C. livia* and *G. gallus*) and GSE128151 (*A. sagrei*).

---

**Supplemental Figure 1.**
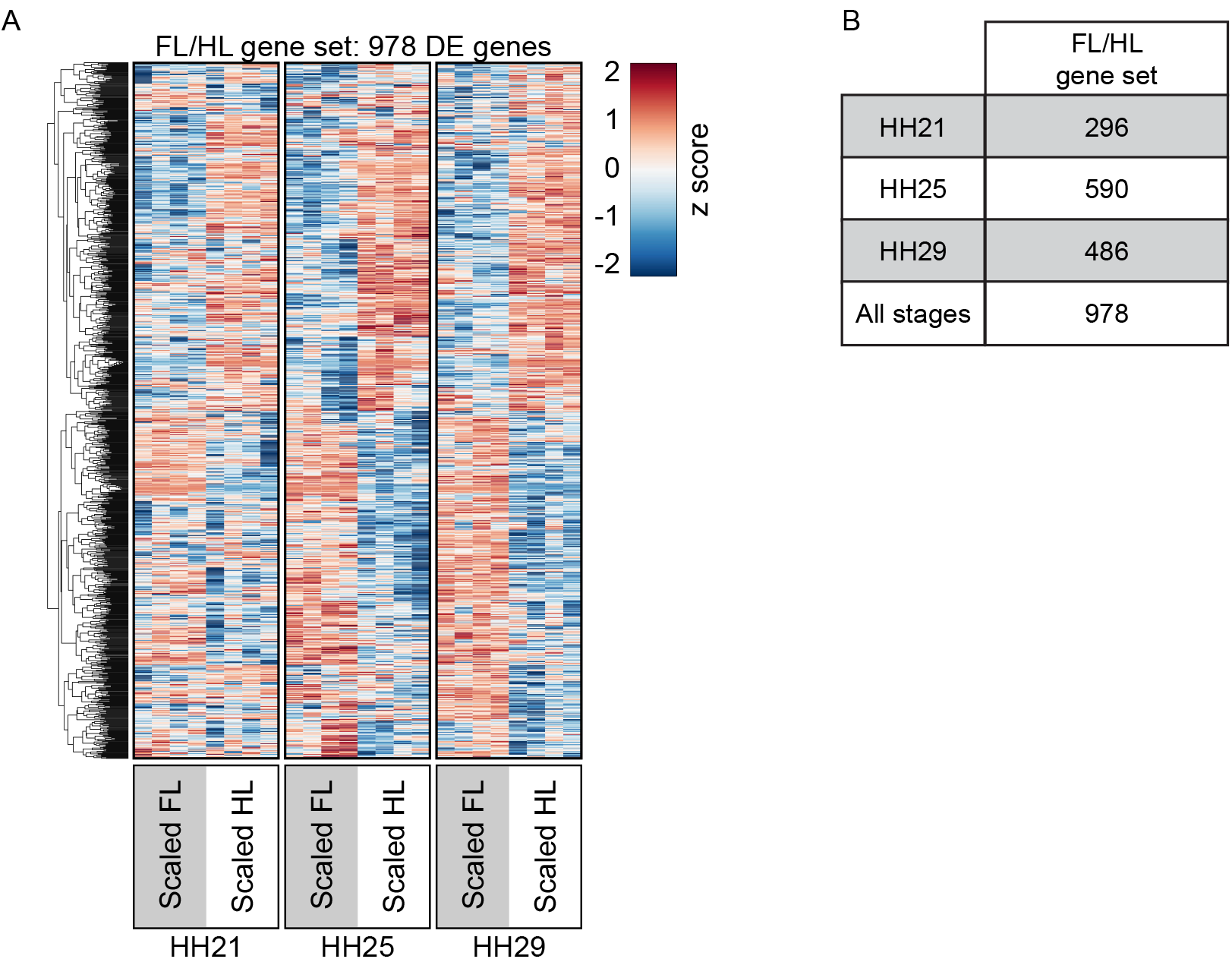
Differentially expressed genes between pigeon FL and scaled HL buds. (**A**) Gene-wise hierarchical clustering heat map showing 978 genes differentially expressed between pigeon FL and scaled HL buds at HH21, HH25, and/or HH29. All genes that are significantly differentially expressed at one or more developmental stage are included in heat map. Z score scale represents mean subtracted regularized log transformed read counts. (**B**) Table summarizing number of significantly differentially expressed genes between FL and scaled HL at HH21, HH25, HH29, or at all stages combined.

**Supplemental Figure 2.**
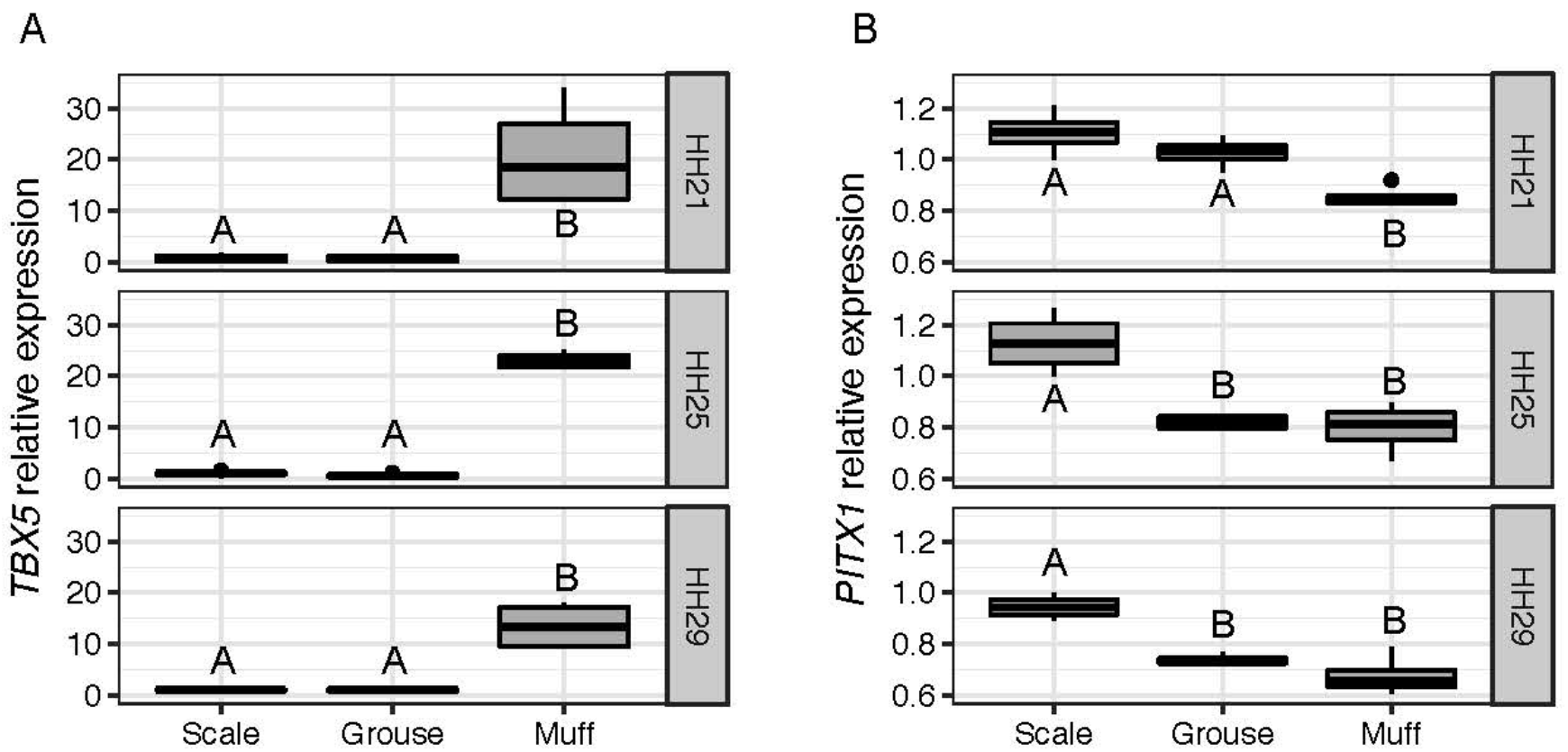
Temporal dynamics of *TBX5* and *PITX1* expression in pigeon limb buds revealed by qRT-PCR. (**A-B**) qRT-PCR analyses of *TBX5* (A) and *PITX1* (B) expression relative to *ACTB* in HH21, HH25, and HH29 pigeon hindlimb buds. Boxes span 1^st^ to 3^rd^ quartiles, bars extend to minimum and maximum values, black line denotes median, dot indicates outlier. For scale n=4 sets of hindlimb buds, grouse n=4, muff n=10. Letter above box (A or B) denotes significance group.

**Supplemental Figure 3.**
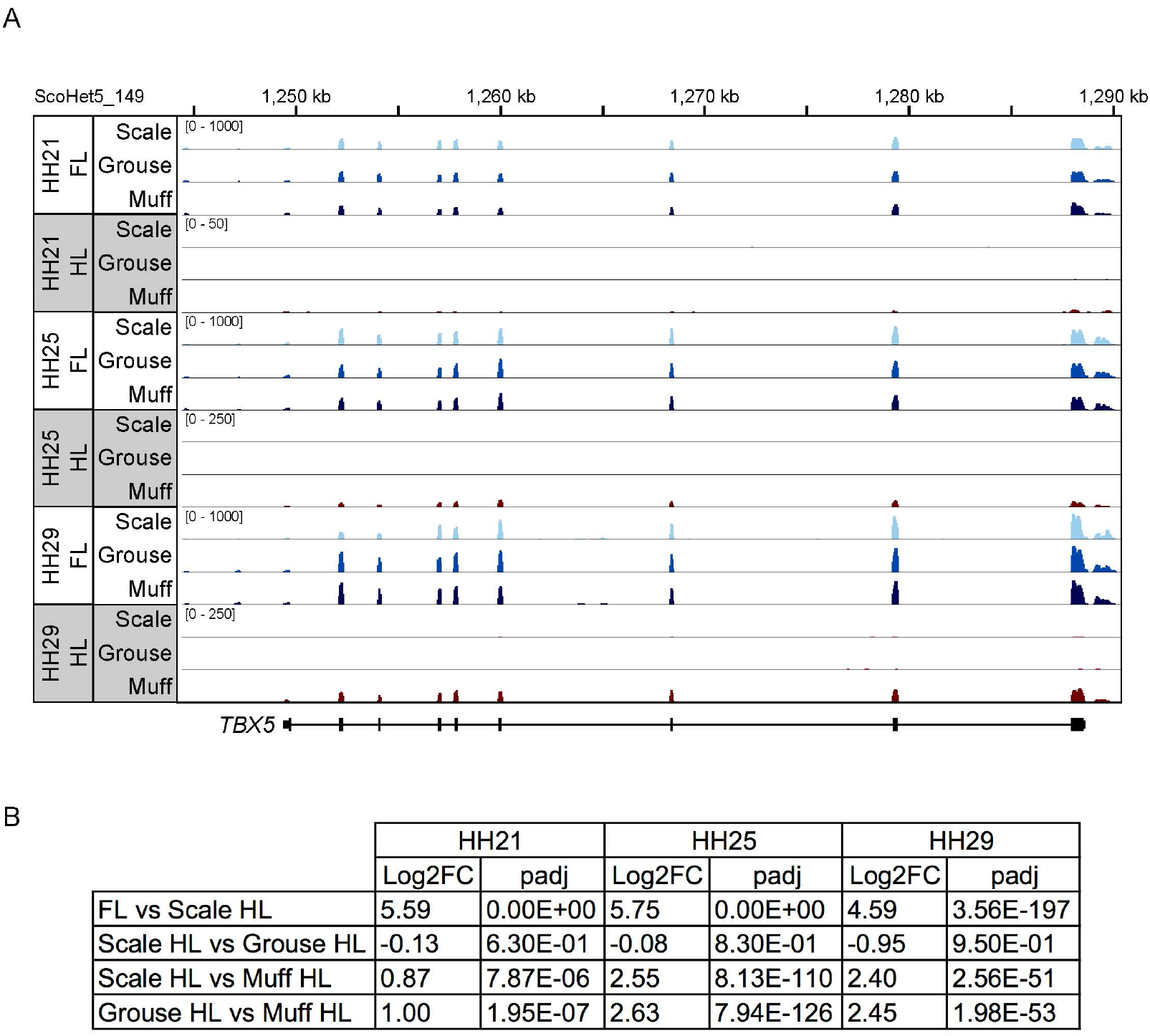
Differential *TBX5* expression in pigeon limb bud samples. (**A**) Representative RNA-seq tracks for *TBX5* in FL and HL bud samples from scaled, grouse, and muff embryos at HH21, HH25, and HH29. Scale is 0-1000 reads for FL tracks and 0-250 reads for HL tracks. (**B**) Table showing *TBX5* log2 fold change (Log2FC) and corresponding adjusted p-value (padj) for each comparison included in differential expression analyses.

**Supplemental Figure 4.**
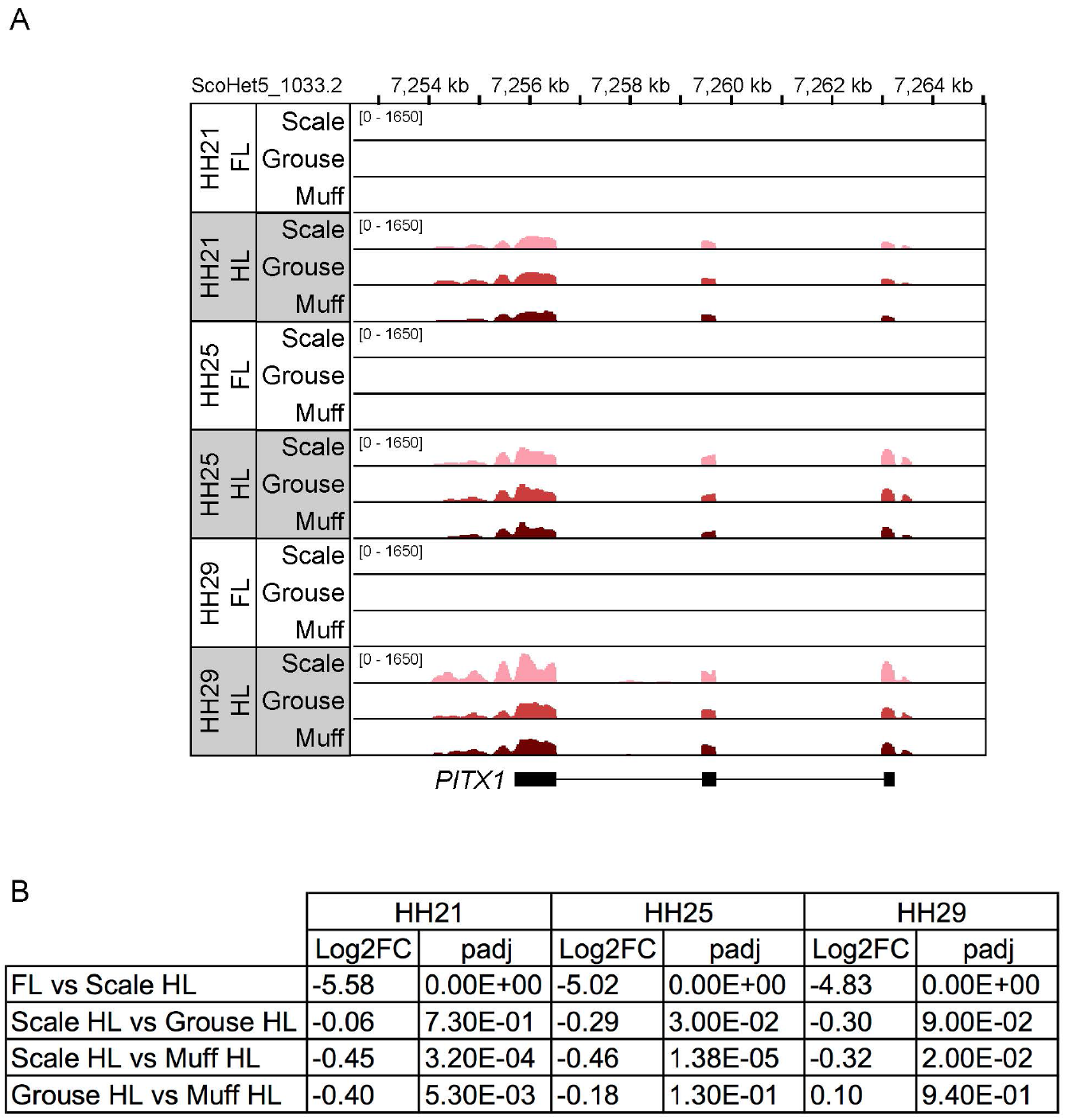
Differential *PITX1* expression in pigeon limb bud samples. (**A**) Representative RNA-seq tracks for *PITX1* in FL and HL bud samples from scaled, grouse, and muff embryos at HH21, HH25, and HH29. Scale is 0-1650 reads for all tracks. (**B**) Table showing *PITX1* log2 fold change (Log2FC) and corresponding adjusted p-value (padj) for each comparison included in differential expression analyses.

**Supplemental Figure 5.**
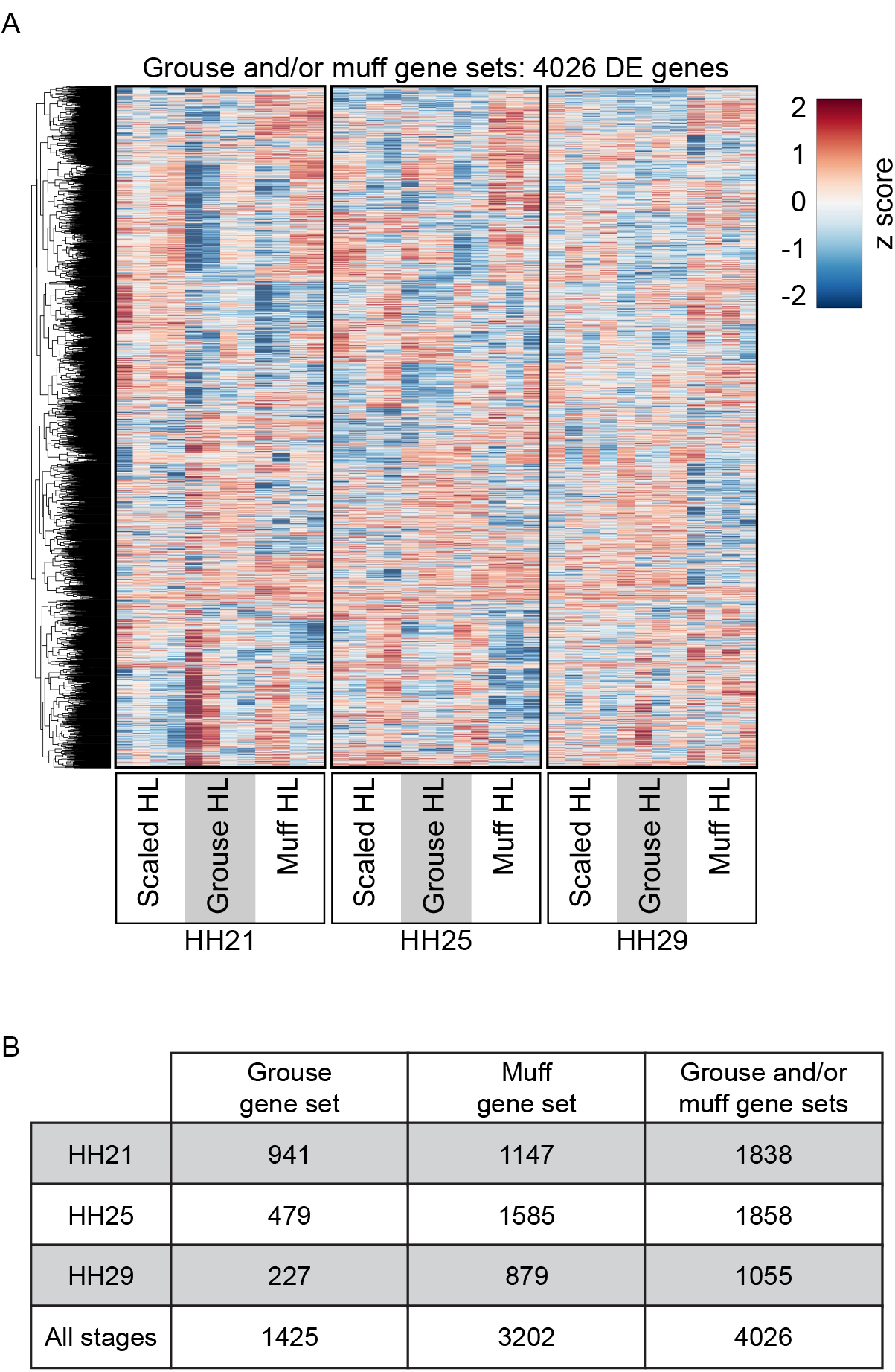
Differentially expressed genes between scaled and feathered pigeon HLs. (**A**) Gene-wise hierarchical clustered heat map showing 4026 genes differentially expressed between HL buds from scaled, grouse, and muff embryos at HH21, HH25, and/or HH29. All genes that are significantly differentially expressed in at least one comparison at one or more developmental stage are included in heat map. Z score scale represents mean subtracted regularized log transformed read counts. (**B**) Table summarizing number of significantly differentially expressed genes between scaled, grouse, and muff HL buds at HH21, HH25, HH29, or at all stages combined.

**Supplemental Figure 6.**
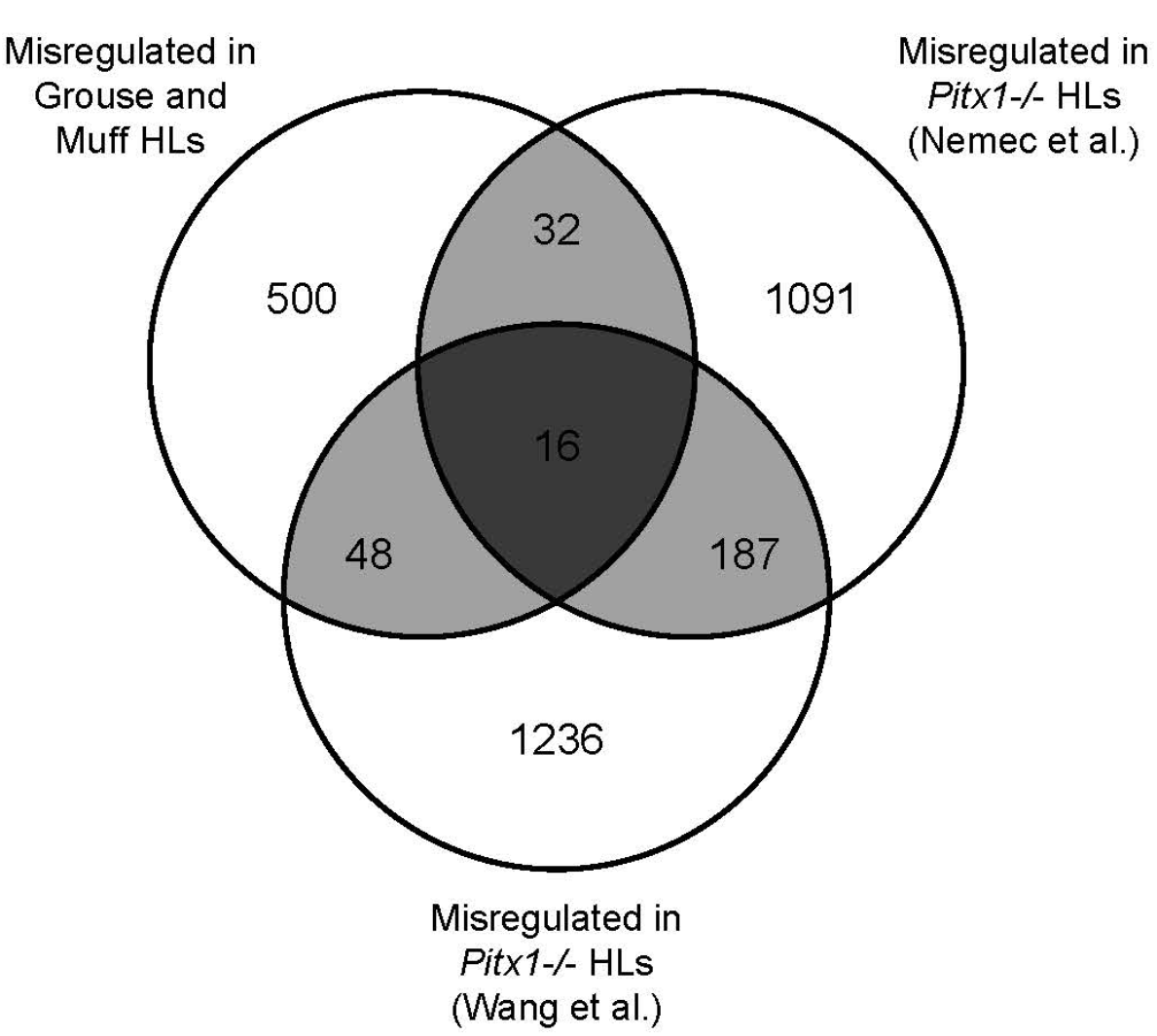
Comparison of *PITX1*-dependent genes in pigeon and mouse limb buds. Venn diagram showing overlap of differentially expressed gene sets from pigeon scaled HL vs. grouse HL and pigeon scaled HL vs. muff HL with two published RNA-seq datasets of mouse wild-type vs. *Pitx1^-/-^* HLs (Nemec et al., 2017, Wang et al., 2018).

**Supplemental Figure 7.**
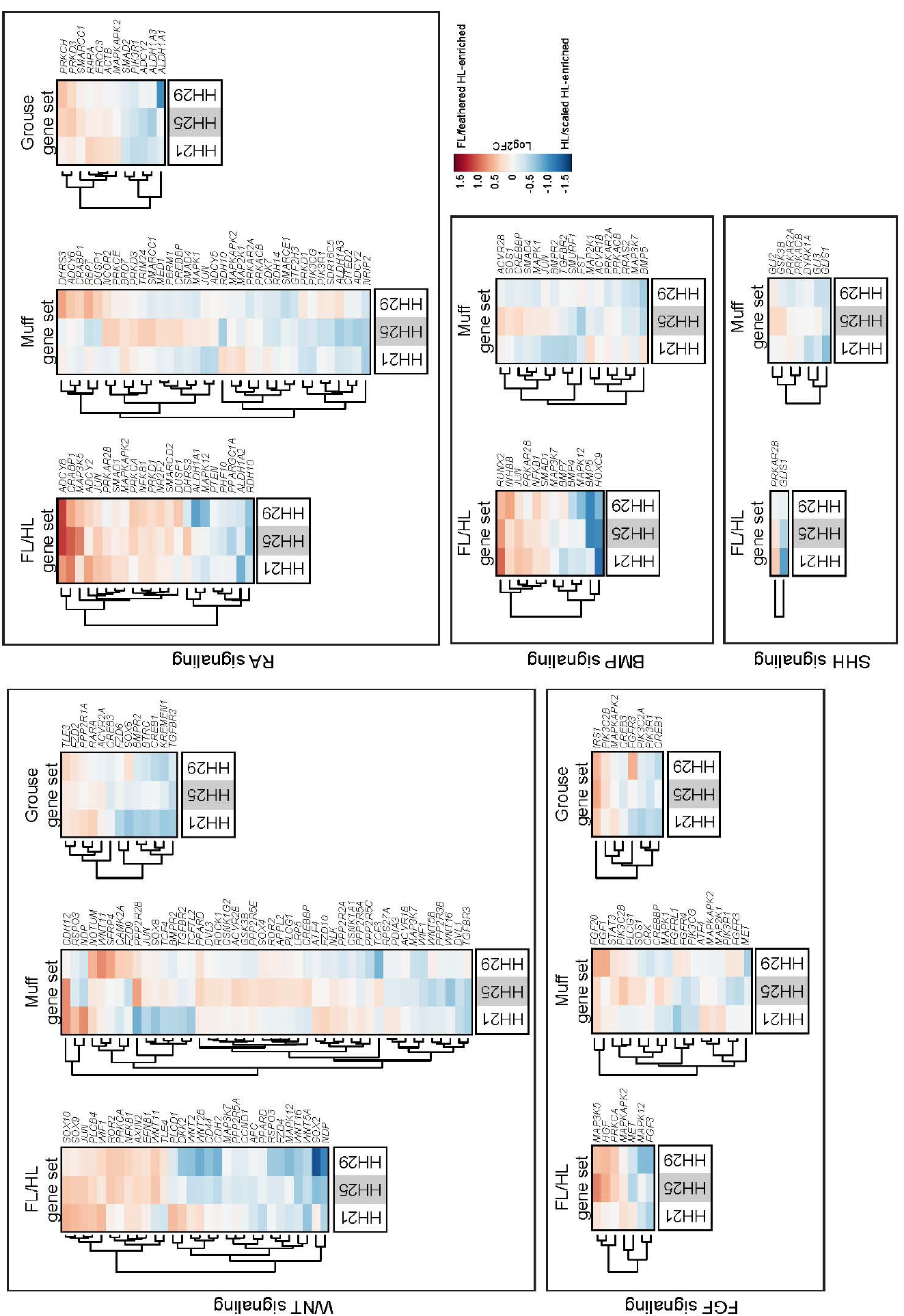
Differentially expressed genes associated with developmental signaling pathways. Gene-wise hierarchical clustering heat maps of differentially expressed genes associated with WNT, FGF, RA, BMP, or SHH signaling pathways. Genes were identified by Ingenuity Pathway Analysis. Scale is Log2FC −1.5 to 1.5.

**Supplemental Figure 8.**
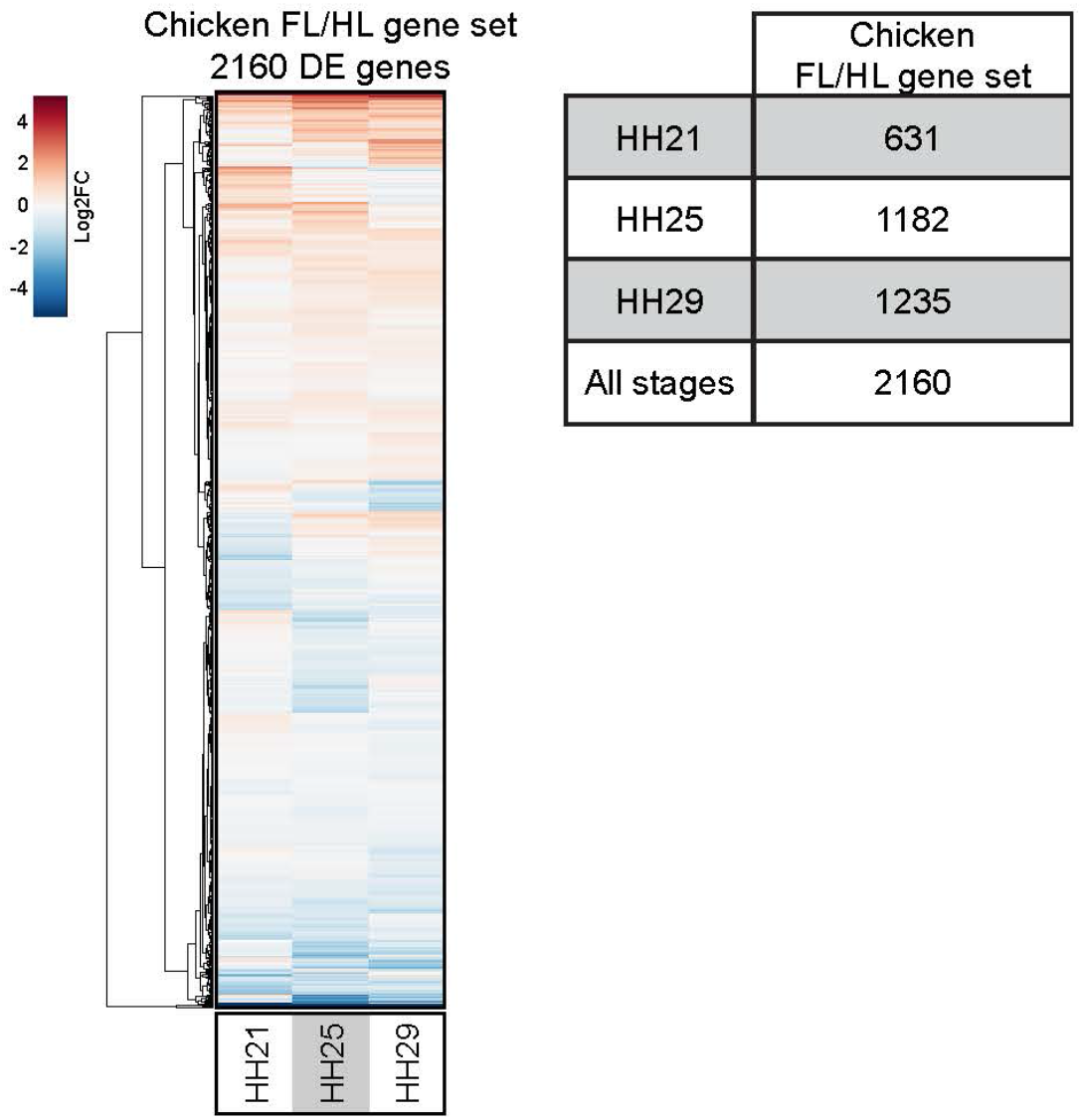
Differentially expressed genes between chicken FL and HL buds. **(A)** Gene-wise hierarchical clustering heat map showing 2160 genes differentially expressed between chicken FL and HL buds at HH21, HH25, and/or HH29. All genes that are significantly differentially expressed at one or more developmental stage are included in heat map. Scale is Log2FC −5 (HL-enriched) to 5 (FL-enriched). **(B)** Table summarizing number of significantly differentially expressed genes between chicken FL and HL at HH21, HH25, HH29, or at all stages combined.

**Supplemental Figure 9.**
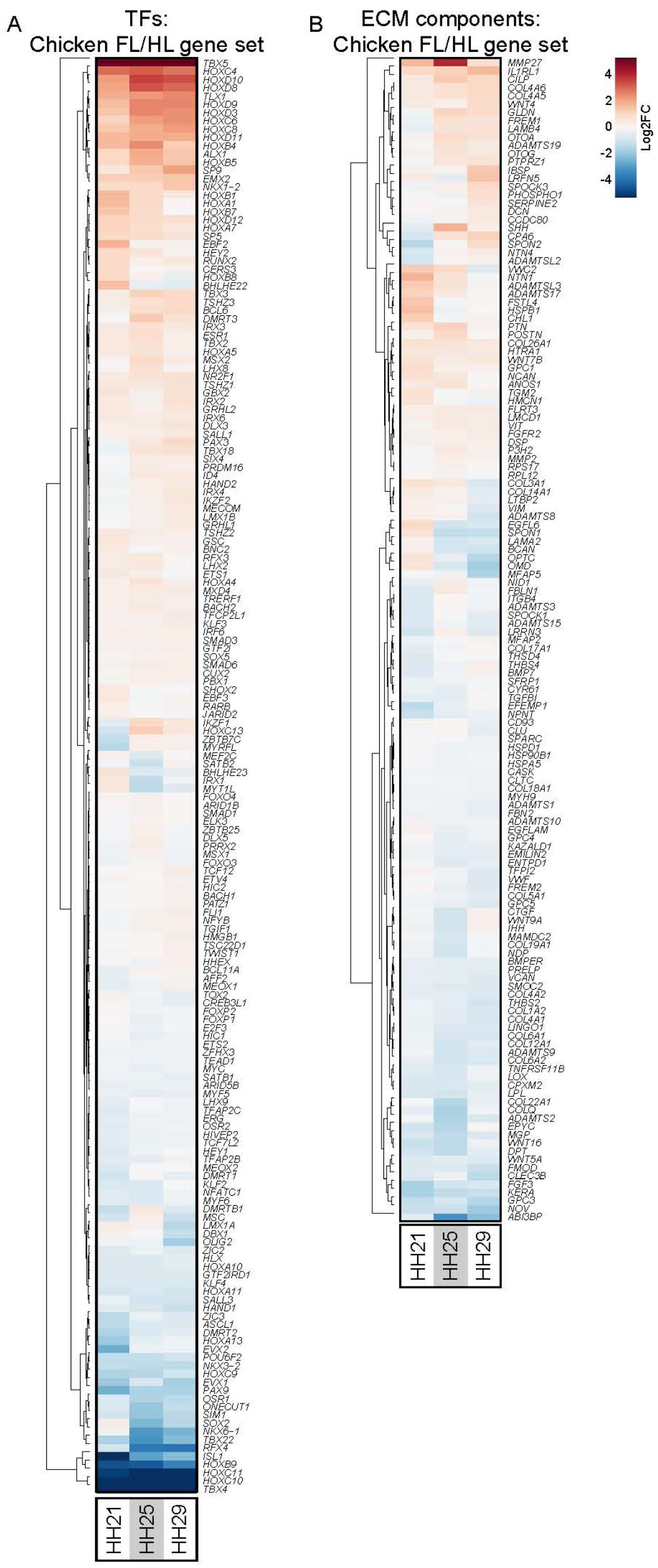
TFs and ECM components differentially expressed between chicken FL and HL. (**A-B**) Hierarchical gene clustering heat maps of all TFs (A) and ECM components (B) that are differentially expressed between chicken FL and HL buds at HH21, HH25, or HH29. Scale is Log2FC −5 (HL-enriched) to 5 (FL-enriched).

**Supplemental Figure 10.**
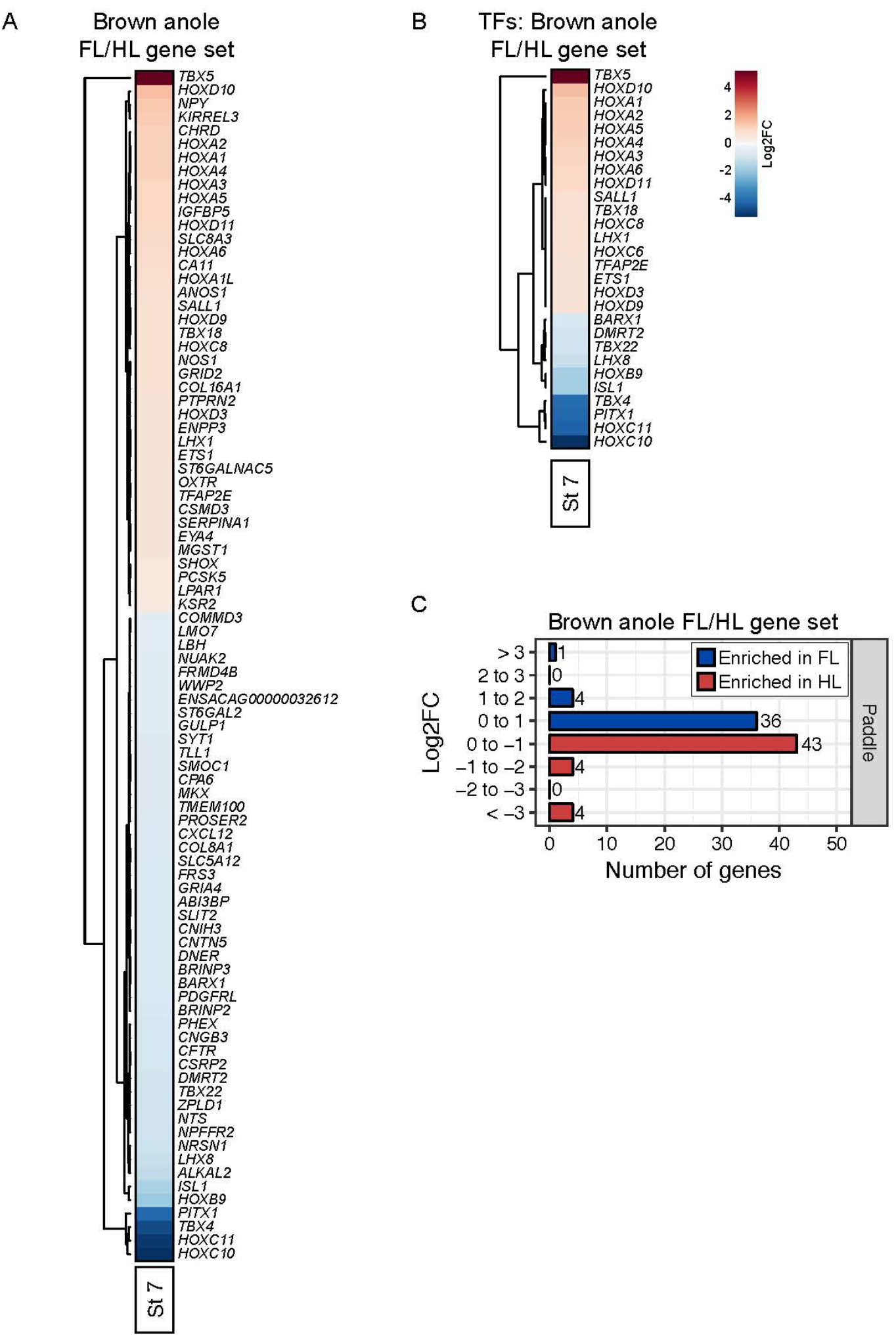
Differentially expressed genes between brown anole FL and HL buds. (**A-B**) Hierarchical gene clustering heat maps of all genes (A) or TFs (B) that are differentially expressed between FL and HL buds from Stage 7 brown anole embryos. Scale is Log2FC −5 (HL-enriched) to 5 (FL-enriched). (**C**) Number of genes enriched in FL or HL specific manner in Stage 7 brown anole embryos. Genes are grouped based on Log2FC from FL vs. HL comparison.

**Supplemental Figure 11.**
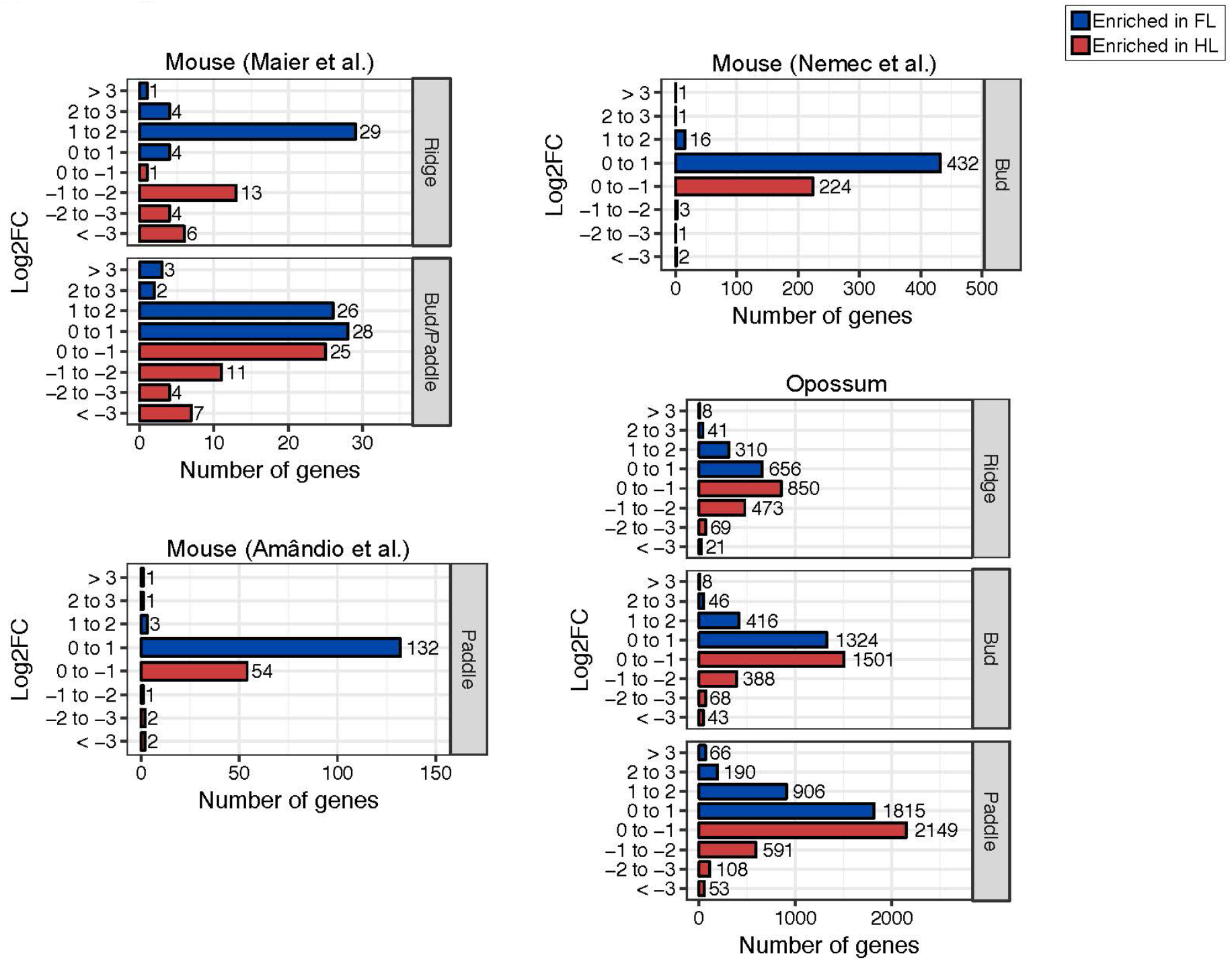
Differentially expressed genes in re-analysis of mammalian FL and HL datasets. (**A-D**) Number of genes enriched in FL or HL specific manner identified in opossum (A), and mouse (B-D) datasets. Genes are grouped based on Log2FC from FL vs. HL comparisons.

